# Canonical Wnt and TGF-β/BMP signaling enhance melanocyte regeneration and suppress invasiveness, migration, and proliferation of melanoma cells

**DOI:** 10.1101/2022.03.11.483949

**Authors:** Esra Katkat, Yeliz Demirci, Guillaume Heger, Doga Karagulle, Irene Papatheodorou, Alvis Brazma, Gunes Ozhan

## Abstract

Tissue regeneration and cancer share remarkable features including activation of cell proliferation and migration. Yet, tumors considerably differ from the regenerating tissue with respect to abnormal proliferation, invasive growth, and metastasis. Thus, it is likely that cancer resembles early stages of regeneration with increased proliferation, but separates from the later stages with reduced proliferation and enhanced differentiation. Here, by exploiting the zebrafish melanocytes that can efficiently regenerate and be induced to undergo malignant melanoma, we unravel the transcriptome profiles of the regenerating melanocytes during early and late regeneration, and the melanocytic nevi and malignant melanoma. Our global comparison of the gene expression profiles of melanocyte regeneration and nevi/melanoma uncovers the differential regulation of a substantial number of genes related to Wnt signaling and TGF-β/BMP signaling pathways between regeneration and cancer. Functional activation of canonical Wnt or TGF-β/BMP pathways during melanocyte regeneration promoted melanocyte regeneration and potently suppressed the invasiveness, migration, and proliferation of human melanoma cells *in vitro* and *in vivo*. Therefore, differential regulation of signaling mechanisms between regeneration and cancer can be exploited to stop tumor growth and develop new anticancer therapies.

## Introduction

The regenerative response of animals to damaged or lost tissues in injury or disease can occur through distinct routes including activation of resident stem cells, the direct proliferation of differentiated cells, and dedifferentiation of cells. These routes all involve the stimulation of cell proliferation, which is also tightly associated with cancer. Cell proliferation has long been proposed to be a major event that links regeneration and tumorigenesis, with the assertion of “individuation fields” both as developing regions where organizers can rearrange the tissues to become a complete embryo and as regulators of tissue growth in the adult organisms (Waddington 1935, Stern 2000). The presence of these fields in adult organisms with high regenerative potential such as urodeles and larval anurans may argue that tissues escaping from the organizing pattern of the fields could grow into cancer (Waddington 1935).

Under normal conditions, regeneration follows a common route of injury-induced events that include cell proliferation, migration, differentiation, and morphogenesis to restore tissue integrity and function (King and Newmark 2012). Most importantly, these events are terminated in a controlled manner to prevent malignant transformation of the tissue (Gurtner, Werner et al. 2008, Eming, Martin et al. 2014, MacCarthy-Morrogh and Martin 2020). However, abnormal progression of regeneration could convert the healing tissue into a rapidly proliferating tumor that fails to establish tissue integrity (Haddow 1972, Coussens and Werb 2002, Beachy, Karhadkar et al. 2004, Gurtner, Werner et al. 2008, Schafer and Werner 2008, Oviedo and Beane 2009). In addition to the elevated levels of cell proliferation, clinical observations such as ulceration and tumor formation during abnormal wound repair, activation of an inflammatory response, angiogenesis, and differential expression of genes involved in cell proliferation, survival, and migration have supported the hypothesis that proposes cancers as non-healing wounds (Dolberg, Hollingsworth et al. 1985, Riss, Khanna et al. 2006, Schafer and Werner 2008, DiPietro 2013, Crusz and Balkwill 2015, Dvorak 2015, DiPietro 2016, Karin and Clevers 2016). These observations thus rationalize a potential link between regeneration and cancer. Nevertheless, the underlying mechanistic link between the two phenomena has been mostly overlooked at the molecular level concerning genome-wide analysis and comparison of gene expression in regenerating tissue and cancer tissue.

Melanocytes constitute a heterogeneous group of melanin-producing cells with diverse roles from pigmenting the skin and providing it with protection against UV damage to neuroendocrine functions (Takeda, Takahashi et al. 2007, Plonka, Passeron et al. 2009). Adult melanocytes belong to the skin, which is the largest and one of the most regenerative organs in vertebrates. Studies on the physiological regeneration of mammalian hair follicles have proven that melanocytes are derived from melanocyte stem cells (MSCs), which are capable of self-renewal and fully competent to produce mature melanocytes (Oshima, Rochat et al. 2001, Nishimura, Jordan et al. 2002, White and Zon 2008). Microphthalmia-associated transcription factor (Mitf), Endothelin receptor B (Ednrb), c-Kit, Notch, and Wnt signaling pathways have been found to play important roles in the maintenance of MSCs and melanocyte regeneration in adulthood (Rawls and Johnson 2001, McGill, Horstmann et al. 2002, Kumano, Masuda et al. 2008, O’Reilly-Pol and Johnson 2013, Iyengar, Kasheta et al. 2015, Guo, Xing et al. 2016, Takeo, Lee et al. 2016, Watanabe, Motohashi et al. 2016). The biological processes and pathways including Mitf, Ednrb, c-Kit, and Notch that are characteristic of melanocyte regeneration have also been associated with melanoma, the most aggressive and deadliest form of skin cancer originating from the melanocyte lineage (Sulaimon and Kitchell 2003, Widlund and Fisher 2003, White and Zon 2008, Krauss, Frohnhöfer et al. 2014, Liu, Fukunaga-Kalabis et al. 2014, Wang, Liu et al. 2018). Here, we postulate that the molecular mechanisms of tumorigenesis should show more parallelism with those of regeneration at its relatively earlier stages. Yet, the mechanisms could differ later when regeneration is completed but cancer cells continuously proliferate. Thus, it is rational to assume that while the early-expressed genes are more commonly shared between the two phenomena, late-expressed genes rather differ. To address this assumption, on one side, we took advantage of adult zebrafish melanocytes and examined different stages of regeneration quantitatively for the expression of proliferation and differentiation markers. Accordingly, we determined 1-day post-ablation (dpa) as the “proliferative or early regeneration stage” and 7 dpa as the “differentiation or late regeneration stage”. Comparative genome-wide transcriptome profiling of the regenerating melanocytes in the caudal fin, which we selected as a common platform for comparative analysis of differentially expressed genes (DEGs), revealed that 1 dpa and 7 dpa are transcriptionally distinct stages of melanocyte regeneration. On the other side, we exploited an adult zebrafish model of melanoma, Tg*(mitfa:Hsa.HRAS^G12V^,mitfa:GFP)* line, that enabled the characterization of both melanocytic nevi and malignant melanoma. Transcriptome analysis demonstrated that nevi and melanoma shared DEGs related to melanocyte differentiation and pigmentation but separated from each other by the genes associated with proliferation and neural crest cell (NCC) signature. 1 dpa and melanoma were mostly enriched for cell cycle and DNA replication signatures, while 7 dpa is hallmarked by the activation of melanocyte differentiation and pigmentation genes. By globally comparing the transcriptomes of the early and late melanocyte regeneration stages to those of nevi and melanoma, we found that Wnt and TGF-β/BMP signaling pathways were differently regulated between melanocyte regeneration and melanoma. Activation of canonical Wnt and TGF-β/BMP signaling enhanced melanocyte regeneration and suppressed the invasiveness, migration, and proliferation of melanoma cells *in vitro* and *in vitro*. Overall, by comparing the stage-dependent alterations in gene expression profiles between melanocyte regeneration and melanoma, our study pioneers testing the potential of interfering with cancer progression.

## Materials and Methods

### Zebrafish husbandry and maintenance

Zebrafish are maintained following the guidelines of the Izmir Biomedicine and Genome Center’s Animal Care and Use Committee. All animal experiments were performed with the approval of the Animal Experiments Local Ethics Committee of Izmir Biomedicine and Genome Center (IBG-AELEC).

### Adult zebrafish melanocyte ablation and subsequent regeneration

The zebrafish melanocytes were ablated with Neocuproine (NCP; Sigma-Aldrich, MO, USA), a copper chelating agent that specifically kills mature melanocytes without giving harm to *mitf+* melanocyte stem/progenitor cells (O’Reilly-Pol and Johnson 2008, Iyengar, Kasheta et al. 2015). 6-12 month-old wild-type (wt) AB zebrafish were treated with 1 µM of NCP for 24 hours and caudal fins were collected at 1, 2, 3, 4, 7, 15, and 20 days post-ablation (dpa) (Figure 1, left; Figure S1A).

**Figure 1.**
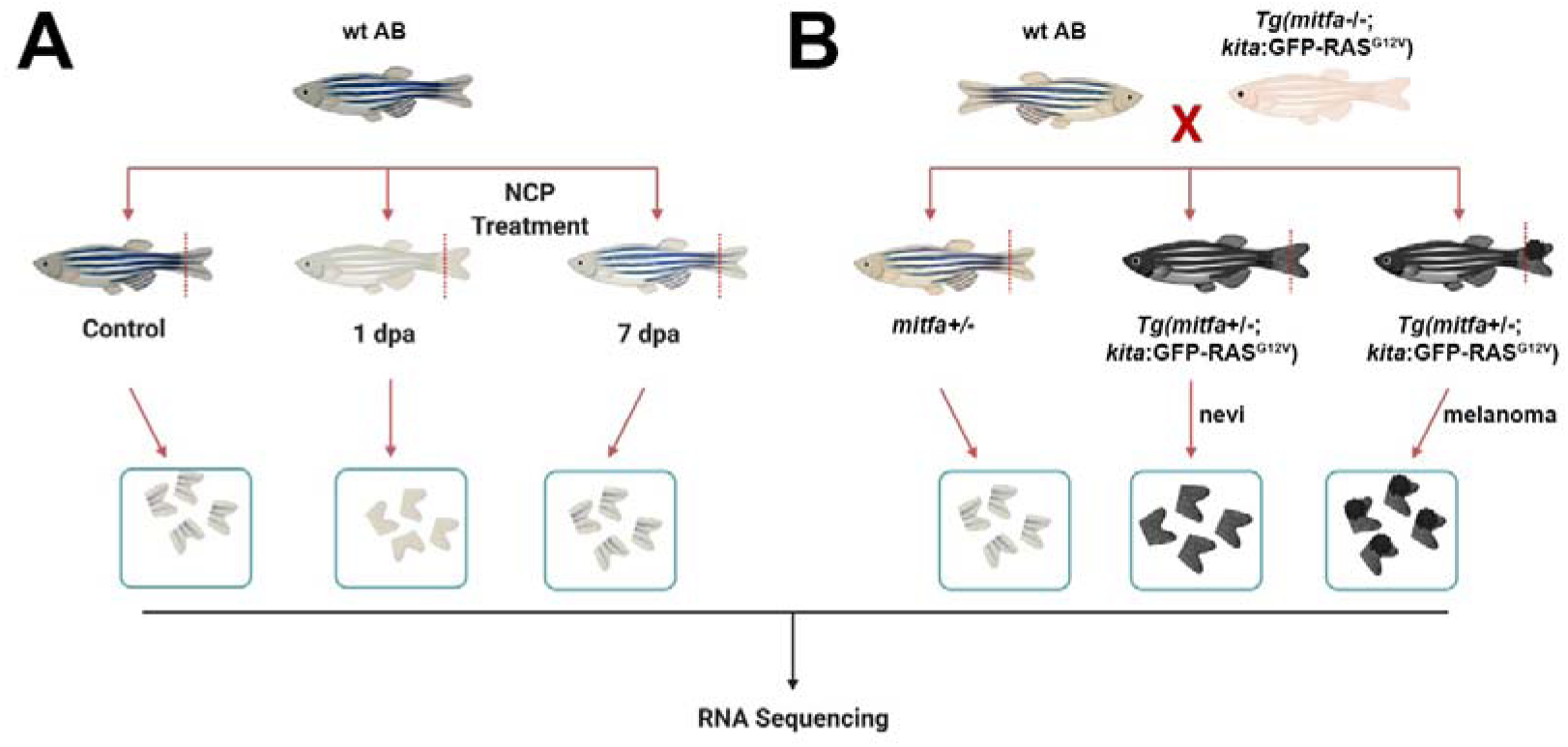
Generation of the zebrafish models of melanocyte regeneration and melanoma. **(A)** Following NCP treatment, caudal fins of individual zebrafish were collected for each group (control, 1 dpa, and 7 dpa) and used as biological replicates (no pooling). **(B)** The zebrafish melanoma model was generated by outcrossing the Tg*(mitfa:Hsa.HRAS^G12V^,mitfa:GFP)* line with the wt AB zebrafish. 50% of the siblings (*mitfa*+/-) were used as the control group. 50% of the siblings Tg *(mitfa+/-,HRAS^G12V^:GFP)* were used for the collection of the nevi and melanoma tissues. Caudal fins were collected for each group and used as biological replicates (no pooling).

Samples from 1, 2, 3, and 4 dpa and samples from 7, 15, and 20 dpa were assayed for early and late melanocyte regeneration, respectively. First, fish were anesthetized with 10 mg/mL of Tricaine (MilliporeSigma, MA, USA). Following immobilization, the caudal fin of each fish was resected to 50% of its original size along the dorsoventral axis with a razor blade and stored in RNAlater until RNA isolation. Four animal fins were used for each regeneration stage.

### Adult zebrafish model of melanoma

The zebrafish melanoma was obtained by using Tg*(mitfa:Hsa.HRAS^G12V^,mitfa:GFP)* line as described previously (Michailidou, Jones et al. 2009). First, this zebrafish line was outcrossed to the wt (AB) zebrafish to generate 50% Tg*(mitfa+/-;mitfa:Hsa.HRAS^G12V^,mitfa:GFP)* embryos that could grow melanocytes due to the single functional copy of *mitfa* and expressed *RAS^G12V^* oncogene only in the melanocytes due to the *mitfa* promoter, leading to hyperpigmentation throughout the body (Figure 1, right). The remaining 50% of embryos were Tg(*mitfa+/-*) and used as control.

### RNA isolation and quantitative PCR

RNA isolation from the fin tissues was carried out using the RNeasy® micro kit (Qiagen, Germany), and from SKMEL-28 cells with QIAzol lysis reagent (Qiagen, Germany), according to the manufacturer’s instructions. The fin tissue was homogenized with RNase-free disposable pestles and the lysate was passed through a 20G insulin needle 5-10 times. The samples were treated with DNAse I (Qiagen, Germany) to prevent genomic DNA contamination. Total RNA concentration was determined in a NanoDrop™ 2000/C spectrophotometer (Thermo Fisher Scientific, MA, USA). RNA quality was verified using RNA 6000 Pico kit (Agilent Technologies, CA, USA) according to the manufacturer’s instructions, and samples were validated to have RNA integrity values above 7.0. cDNA synthesis was performed by using the iScript^TM^ cDNA synthesis kit (Bio-Rad Laboratories, CA, USA). qPCR was performed in triplicates by using GoTaq qPCR Master Mix (Promega, WI, USA) in an Applied Biosystems 7500 Fast Real-Time PCR machine (Thermo Fisher Scientific, MA, USA). For zebrafish samples, expression values were normalized to *Danio rerio ribosomal protein L13a (rpl13a)*, the reference housekeeping gene. For melanoma cell line samples, expression values were normalized to human *Glyceraldehyde 3-phosphate dehydrogenase GAPDH*, the reference housekeeping gene. Primer sequences are provided in Table S1.

### Library construction and RNA sequencing (RNA-seq)

The samples with an RNA Integrity Number (RIN) ≥ 8 were selected for RNA-seq. RNA quality was also checked by performing quantitative reverse transcription PCR (RT-PCR) with a primer pair generating an 812-bp product for zebrafish *beta-actin 1* (*actb1*). The samples that passed the quality controls were sent to the Genomics Core Facility (GeneCore, EMBL, Germany) for library preparation and RNA-seq. An Illumina TruSeq RNA Library Preparation Kit v2 (Illumina, CA, USA) was used to prepare libraries according to the instructions given by the manufacturer. For each reaction, 500 ng of cDNA was used. A paired-end, strand-specific sequencing platform was used on Illumina NextSeq 500 (Illumina, CA, USA) with a read length of 75 bp.

### Bioinformatics analysis

Read quality control of RNA-seq samples obtained from the zebrafish melanocyte regeneration stages (1 dpa and 7 dpa), zebrafish melanocytic nevi and zebrafish malignant melanoma was initially performed by using the FastQC tool (Andrews 2010). The reads were aligned to the zebrafish reference genome GRCz11 (danRer11) by using HISAT2 (version 2.1.0) (Kim, Langmead et al. 2015)_. The read summarizer HTSeq 0.6.0_ was used to count transcripts by using the annotation file Danio_rerio.GRCz11.93.gtf from the Ensembl database. Gene expression counts were normalized and tested for differential expression using DESeq2 (Love, Huber et al. 2014). All genes were tested for differential expression at the two stages of melanocyte regeneration (1 dpa and 7 dpa), in nevi and melanoma, all in contrast to the control baseline zebrafish transcriptome, using a Wald test on log2 fold changes. In all comparisons, differentially expressed genes (DEGs) were called with a false discovery rate (FDR) of 0.1 and an effect size threshold of log2(1.2) (thereafter referred to as FC>1.2 in either direction). Gene ontology (GO) and Kyoto Encyclopedia of Genes and Genomes (KEGG) pathway enrichment analyses were carried out with lists of all DEGs against the DAVID knowledgebase (Huang da, Sherman et al. 2009). The EASE score, a modified one-tailed Fisher’s exact test, was used to measure the significance of the terms’ and pathways’ enrichment. Gene Set Enrichment Analysis (GSEA) was performed in pre-ranked mode using the fgsea Bioconductor package (Subramanian, Tamayo et al. 2005, Korotkevich, Sukhov et al. 2021). KEGG pathways collection was retrieved via the KEGGREST package (Tenenbaum and Maintainer 2022). Ensembl IDs of all zebrafish genes were converted into Entrez IDs by using the AnnotationDbi package (Pagès, Carlson et al. 2022). Genes were ranked using the Wald statistic computed by DESeq2. Significant gene sets were selected according to an FDR < 0.05. The data was visualized by principal component analysis (PCA) using the packages ggplot2 (version 3.3.2) and pheatmap (version 1.0.12) (Kolde 2012, Wickham 2016). Gene lists were obtained from the AmiGO (Carbon et al., 2009) and KEGG databases for selected GO terms and pathways. The enrichment of DEGs in GO terms and KEGG pathways were visualized using GOChord plots (Walter, Sanchez-Cabo et al. 2015).

### Drug administration

A stock solution of NCP was prepared as 25 mg/ml in DMSO. The final concentration of NCP was 1 µM. A stock solution of epinephrine (Sigma-Aldrich, MO, USA) was prepared as 18 mg/ml (1 M) in 0.125 M HCl. Before imaging, zebrafish were treated with epinephrine for 15-20 min at a final concentration of 1 mg/ml. The drugs used for the embryo/larva regeneration assay, xenotransplantation, and *in vitro* assays on SK-MEL-28 cells were as follows: 4-Hydroxyanisole (4-HA, Sigma-Aldrich, MO, USA), the canonical Wnt pathway antagonist IWR-1 (Sigma-Aldrich, MO, USA), the GSK-3 inhibitor, and Wnt agonist BIO (Sigma-Aldrich, MO, USA), the selective bone morphogenetic protein (BMP) antagonist DMH-1 (Bio-Vision Inc., CA, USA) and the BMP agonist isoliquiritigenin (ISL; Raybiotech, GA, USA). DMSO was used as a control in drug treatment experiments. Toxicity tests were conducted on zebrafish embryos that were treated with different concentrations of the drugs from 24 hours post-fertilization (hpf) to 7 days post-fertilization (dpf). Based on a survival rate above 90%, the working concentrations of drugs were determined as follows: 10 µM for IWR-1, 1 µM for BIO, 20 µM for ISL, and 5 µM for DMH-1.

### Melanocyte regeneration measurement on zebrafish embryos/larvae

Dechorionated wt (AB) zebrafish embryos were treated with 20 µM 4-HA in embryo water between 36 hpf and 60 hpf to ablate melanocytes and induce melanocyte regeneration. 4-HA is a phenolic compound that is converted to a toxic form of o-quinine in melanocytes, eventually leading to their death (Yang and Johnson 2006). 4-HA-treated larvae at 2 dpf were then incubated with the drugs and were aligned in a 24-well plate (10 larvae/well, 60 larvae/group). Melanocyte regeneration was quantified daily until 7 dpf. Drugs were replenished every other day.

### Wound healing assay

SK-MEL-28 cells were seeded in a 12-well cell culture plate and allowed to adhere overnight. The culture media was removed, and the cells were washed once with phosphate-buffered saline (PBS). Fresh DMEM with 1 μM BIO or 10 μM ISL was added to the wells. A scratch/wound was created in the cell monolayer using a 200 ul pipette tip. Images of the scratch/wound were taken at the beginning of the experiment (0 h) using an inverted phase contrast microscope. The cells were then incubated at 37°C with 5% CO_2_. After 16 hours, images of the same scratch/wound were captured. The gap area was measured at both time points (0 h and 16 h) using ImageJ analysis software using the plugin for the high throughput image analysis of *in vitro* scratch wound healing assay. The wound closure percentage was compared between the control group and the treatment groups to evaluate the drug’s effect on wound healing.

### Phalloidin staining

SK-MEL-28 cells were treated with 1 μM BIO or 10 μM ISL for 12 hours. To detect actin stress fiber formation, the cells were fixed and stained with Alexa Fluor™ 488 Phalloidin (Invitrogen, MA, USA), and imaged using fluorescence confocal microscopy.

### Preparation of cells for xenotransplantation

The human melanoma cells SK-MEL-28 (HTB-72™) were cultured in Dulbecco’s modified eagle medium (DMEM) supplemented with 10% fetal bovine serum (FBS) and 1% Pen/Strep at 37°C in 5% (v/v) CO_2_ humidified environment. On the day of injection, approximately 1.5 x 10^6^ cells were harvested at 70-80% confluence and washed with Dulbecco’s phosphate-buffered saline (DPBS) containing 10% FBS. Next, cells were resuspended in 50 µl of DPBS with 10% FBS and incubated with 2.5 µl of Vybrant® DiI cell-labeling solution (Invitrogen, MA, USA) for 20 min. Cells were then washed once with FBS and twice with DPBS containing 10% FBS. The pellet was then resuspended in DMEM with 10% FBS to a final density of 30.000 cells/µl. Cells were mixed with 1 µl of Phenol Red (Sigma-Aldrich, MO, USA). Cells that were used for immunofluorescence detection were labeled with CellTracker™ Red CMTPX Dye (C34552, Invitrogen, MA, USA) that was compatible with the fixatives.

### Zebrafish larval xenografts and migration assay

2 dpf *casper* (*roy -/-; nacre-/-)* zebrafish larvae were dechorionated with 0.1 mg/mL pronase (Sigma-Aldrich, MO, USA) solution for 10 min at 28°C. Larvae were then anesthetized with 1 mg/mL Tricaine in E3 medium and transferred to a microinjection plate prepared with 3% agarose in E3. Borosilicate glass capillaries (4 inches, OD 1.0 mm, World Precision Instruments, FL, USA) were used for injection. 400-500 cells were injected directly into the yolk sac of the larva. Injections were gently performed into the middle of the yolk sac to avoid any damage to the duct of Cuvier. Larvae were incubated at 34°C in fresh E3 overnight. The next day, larvae showing infiltration of tumor cells into blood circulation were discarded and the remaining larvae were randomly distributed to each experimental group and exposed to the drugs. At 5 days post-injection (dpi), xenografts were imaged using a fluorescence stereomicroscope, and the percentage of micrometastasis was determined.

### In vivo imaging and quantification of xenografts

Zebrafish larvae were anesthetized with 1 mg/ml Tricaine and transferred to a 6-well glass-bottom plate containing 3% methylcellulose (Sigma-Aldrich, MO, USA) in E3 medium. All bright-field and fluorescent images of the larvae were captured with an Olympus SZX2-ILLB stereomicroscope (Olympus Corporation, Japan). Image processing and quantifications were done in FIJI/ImageJ software as described (Martinez-Lopez, Póvoa et al. 2021).

### Whole mount immunofluorescence staining of zebrafish larvae

The immunofluorescence staining procedure previously described by Martinez-Lopez et al. was performed with several modifications (Martinez-Lopez, Póvoa et al. 2021). Larvae were fixed in 4% paraformaldehyde (PFA) in 1X PBS overnight at 4 °C. The next day fixed larvae were washed with 1x PDT (1x PBST, 0.3% Triton-X, 1% DMSO), permeabilized with ice-cold acetone. Larvae were blocked for 2 hours in PBDX GS blocking buffer (%10 bovine serum albumin, %1 DMSO, 0.3% Triton-X, 15 µL/1 mL goat serum) and incubated with the primary antibody in PBS/0.1% Triton at 4 °C overnight. The next day, the larvae were rinsed with PBS/0.1% Triton, incubated with the secondary antibody at RT for 2 hours, refixed in 4% PFA at RT for 20 minutes, and washed with PBS/0.1% Triton. Larvae were mounted in 80% glycerol between two coverslips and stored at 4°C. The primary antibodies were rabbit anti-cleaved-caspase-3 (1:200, 5A1E, Cell Signaling Technology, MA, USA) and rabbit anti-phospho-histone H3 (1:200, 9701, Cell Signaling Technology, MA, USA). The secondary antibodies were Fluorescein (FITC) AffiniPure donkey anti-rabbit IgG (1:200, 711-096-152, Jackson Immunoresearch Laboratories, PA, USA) and Cy5 AffiniPure donkey anti-mouse IgG (1:200, 715-175-150, Jackson Immunoresearch Laboratories, PA, USA). Nuclear staining was carried out by using 4′,6-diamidino-2-phenylindole (DAPI; 4083S, Cell Signaling Technology, MA, USA). Larvae were imaged using fluorescence confocal microscopy.

### Confocal imaging and quantification

Confocal images were recorded with a 25x or a 63X objective lens using the z-stack function with an interval of 10 µm between each slice. Image processing and quantifications were performed using FIJI/ImageJ software by following the previously published protocol (Martinez-Lopez et al., 2021). Briefly, mitotic figures were counted using the counter plugin and divided by the number of DiO+, DAPI+ nuclei in each corresponding slice. To quantify cleaved-caspase-3, all slices were automatically counted on the ROI (Analyze>Analyze Particles) and analyzed as described previously (Martinez-Lopez et al., 2021).

### Western blotting

The larval tissues or cultured cells were homogenized in RIPA lysis and extraction buffer, and centrifuged at maximum speed for 30 minutes at 4°C. The supernatants were collected and mixed with 5X SDS loading buffer. For western blotting, the samples were separated by a 12% acrylamide-bis acrylamide gel and transferred to a nitrocellulose membrane. Blocking was performed with either 5% milk powder or %5 BSA for 1 hour at room temperature, and the samples were incubated with the following primary antibodies overnight at 4°C: Rabbit anti-p44/42 MAPK (Erk1/2) (1:1000, 4695, Cell Signaling Technology, MA, USA), rabbit anti-phospho-p44/42 MAPK (Erk1/2) (Thr202/Tyr204) (1:500, 4370, Cell Signaling Technology, MA, USA), mouse anti-beta Catenin (1:1000, ab22656, Abcam, Cambridge, UK), rabbit anti-phospho-β-Catenin (Ser675) (1:1000, 4176, Cell Signaling Technology, MA, USA), rabbit anti-β-Actin (1:1000, 4967, Cell Signaling Technology, MA, USA), mouse anti-vimentin (1:1000, sc-373717, Santa Cruz Biotechnology, Inc. TX, USA). Secondary antibodies were donkey anti-rabbit IgG HRP-linked LICOR IRDye 800CW (1:2000, LI-COR Biosciences, NE, USA) and goat anti-rabbit IgG DyLight™ 800 4X PEG Conjugate (1:2000 5151 Cell Signaling Technology, MA, USA).

### Larval melanin content assay

Larvae were anesthetized with 10 mg/mL of Tricaine and transferred to 1.5_ml Eppendorf. Larvae were completely dissolved with agitation in 200_μl of 1 M of NaOH at 37_°C in Eppendorf tubes. Then, samples were transferred to a round (U) bottom 96-well plate (20 larvae/well) and absorptions were measured at 340 nm. Casper larvae were used as blank to subtract the signal of the eyes (Fernandez Del Ama, Jones et al. 2016).

### Statistical analysis

The data were analyzed statistically with the GraphPad Prism 9 software (Graphpad Software Inc., CA, USA). Two-tailed Student’s t-test was used for qPCR and analysis of the percentage of micrometastasis. One-way ANOVA and Tukey’s multiple comparison tests were used to compare more than two groups. Statistical significance was evaluated as follows: *p_≤_0.05, **p_≤_0.01, ***p_≤_0.001, ****p_≤_0.0001, and ns: non-significant.

## Results

### Validation of early and late stages of melanocyte regeneration in the adult zebrafish fin at the transcriptional level

The transcriptional dynamics underlying melanocyte lineage progression in the hair bulb have recently been investigated concerning the activation of quiescent MSCs to proliferate and differentiate into mature melanocytes in a mouse model (Infarinato, Stewart et al. 2020). Pigment cells including the melanocytes and iridophores have been analyzed at the transcriptional level (Higdon, Mitra et al. 2013). However, gene expression profiles of the regenerating melanocytes have not been identified at proliferation and differentiation stages so far. Thus, we set out to unravel the DEGs at the early regeneration stage, where proliferation is the prominent event, and the late regeneration stage, where the proliferative response is replaced by differentiation. Initially, to determine the early and late stages of melanocyte regeneration, we ablated melanocytes of the adult zebrafish by 1-day of NCP treatment and monitored NCP-treated zebrafish daily to track the melanocyte death, depigmentation, onset of melanocyte differentiation and reconstitution of the stripe pattern after washout (Figures 2A, S1A). We followed the change in the morphology of the melanocytes until most of the melanin content was disposed of in the skin. Melanocytic aggregates began to disappear at 3 dpa and were substantially lost at 4 dpa (Figures 2A, S1A). It is important to note that, despite losing their dendritic morphologies, some melanocytic aggregates remained as black smears. Thus, we assumed that most of the melanocytes lost their cellular activity at 4 dpa. Pigmentation started all over the body at 7 dpa and the stripe pattern was reconstituted within 30 days after the fish were returned to freshwater (Figures 2A, S1A). We treated the zebrafish with epinephrine, which leads to the assembly of melanin-containing melanosomes towards the nucleus in the presence of an epinephrine signal when they are alive (Johnson, Africa et al. 1995, O’Reilly-Pol and Johnson 2008, Iyengar, Kasheta et al. 2015, Kang, Karra et al. 2015). Whereas the dendritic morphology of melanocytes turned into compact rounds in untreated fish within 15-20 min, melanocytes of NCP-treated fish did not respond at 1 dpa and 2 dpa, confirming the successful ablation of the melanocytes (Figure S1B).

**Figure 2.**
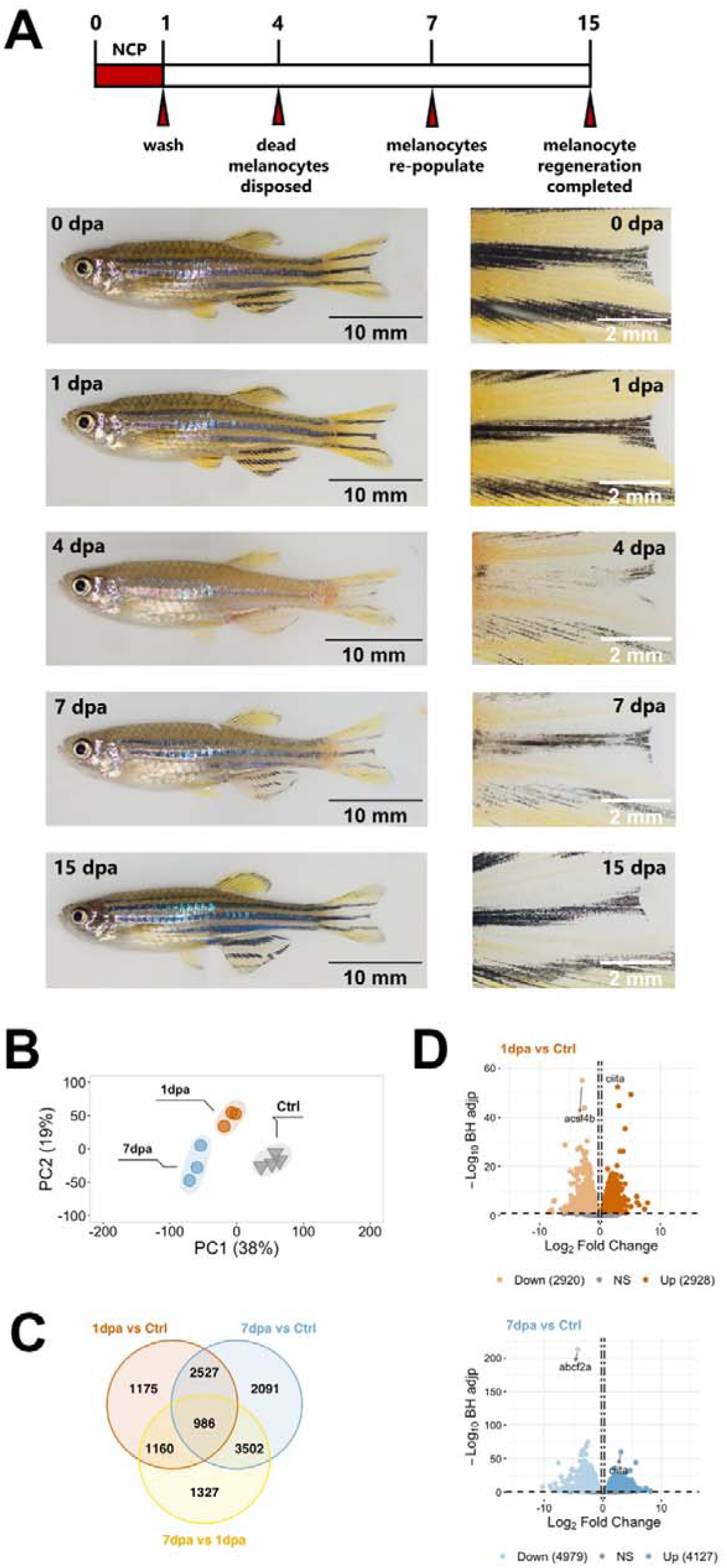
Validation of the melanocyte regeneration model and quantitative analysis of the DEGs at the proliferation and differentiation stages. **(A)** Zebrafish melanocyte regeneration within 15 days after 1 day of NCP treatment. Control zebrafish with normal stripe patterns on the body including the caudal fin before NCP treatment (0 dpa). Melanocytes were lost gradually starting at 1 dpa and extruded out of the skin at 4 dpa when the stripe pattern was not visible anymore. Melanocytes started to become visible at 7 dpa. At 15 dpa, melanocyte regeneration was completed and the stripe pattern was re-established. Scale bars are 5 mm (whole fish on the left) and 2 mm (close-ups on the right). **(B)** Principal component analysis (PCA) of proliferative (1 dpa) and differentiation (7 dpa) stages of melanocyte regeneration and their controls. Different colors of dots and triangles represent the sample condition. Three sample groups were well clustered among their replicates and well separated from other sample groups. Principal Component 1 (PC1, x-axis) represents 38% and PC2 (y-axis) represents 19% of the total variation in the data. **(C)** The Venn diagram shows the number of differentially expressed genes (DEGs) in 1 dpa vs control (orange), 7 dpa vs control (blue), 7 dpa vs 1 dpa (yellow), and the overlap between each set of DEGs of melanocyte regeneration. **(D)** Volcano plots represent changes in gene expression levels in 1 dpa and 7 dpa. Each circle represents a gene. The x-axis shows log_2_ of fold change in the condition compared to the control and the vertical dashed lines indicate the fold change cutoffs. The y-axis depicts -log_10_ of the Benjamini-Hochberg adjusted p-value, where the significance threshold is indicated by a dashed horizontal line. Darker and lighter shades indicate upregulation and downregulation, respectively. dpa: days post-ablation.

Next, we aimed to determine the early and late stages of melanocyte regeneration and isolated total RNA from caudal fin tissues collected at 1, 2, 3, 4, 7, 12 and 20 dpa. Tissue samples from four fish were pooled. We used proliferation as the early stage signature of melanocyte regeneration since half of the newly generated melanocytes are differentiated from a set-aside population of *mitfa+* progenitors that undergo rounds of proliferation (Iyengar, Kasheta et al. 2015, Kang, Karra et al. 2015). *Proliferating cell nuclear antigen (pcna)* expression increased 4-fold at 1 dpa, abruptly decreased at 2 dpa, and returned to the control levels at 7 dpa (Figure S2A). Immunofluorescence staining for phospho-histone H3 confirmed that proliferation increased in the adult fin tissue after NCP treatment (Figure S2B). As the early strong activation of proliferation is consistent with the previous data obtained from adult melanocyte regeneration (Iyengar, Kasheta et al. 2015), we determined 1 dpa as the early stage of melanocyte regeneration. For the late stage, we exploited the master regulator of melanocyte development *mitfa* and the late melanocyte differentiation markers *dopachrome tautomerase (dct)* and *tyrosinase (tyr)* (Rawls and Johnson 2001, Ceol, Houvras et al. 2008). The expression levels of *mitfa*, *dct,* and *tyr* started to increase at around 3 dpa, reached their peak at 7 dpa, and inclined toward control levels at 12 dpa (Figure S2A). Taken together with the expression of the proliferation marker at control levels at 7 dpa, we considered 7 dpa as the late melanocyte regeneration stage. This also correlates with our track of melanocyte pigmentation at the macroscopic level where new melanocytes appeared throughout the body at 7 dpa (Figure S1A). The changes in the expression of proliferation and differentiation markers for the individual fin samples were consistent with the pooled fin samples (Figure S2C). Thus, we decided to exploit the zebrafish caudal fin, which successfully confirmed the activation of the marker genes at different stages of regeneration. Next, to identify the differentially expressed genes at the early and late stages of melanocyte regeneration, we collected the caudal fin tissues of the DMSO-treated and NCP-treated zebrafish (Figure 1) and performed RNA-seq. Principal component analysis (PCA) revealed that the samples of 1dpa and 7 dpa were well separated from each other and the control samples (Figure 2B). We have identified 5848 genes (2928 upregulated=Up and 2920 downregulated=Down) and 9106 genes (4127 Up and 4979 Down) that were differentially expressed in response to melanocyte ablation at 1 dpa and 7 dpa, respectively (Figure 2C-D, Table S2). Interestingly, while only 40% of the DEGs detected at 1 dpa were unique to this stage, at 7 dpa the percentage of unique DEGs increased to 61%. Together, these results indicate that 1 dpa and 7 dpa represent stages of proliferation (early) and differentiation (late), respectively, and validate them as two distinct stages of melanocyte regeneration at the transcriptional level.

### Early/proliferative and late/differentiation stages of melanocyte regeneration have distinct transcriptional profiles

To understand how gene expression profiles alter throughout the proliferation and differentiation stages of melanocyte regeneration, we plotted the heatmap of the expression of a group of selected genes that are involved in neural crest cell (NCC) differentiation, epithelial to mesenchymal transition (EMT), melanocyte differentiation/pigmentation, cell cycle, immune response, proliferation and stem cell differentiation (Figure 3A). The heatmap shows the normalized counts for the biological replicates. The baseline expression in control groups is used to determine the direction of regulation at 1 dpa and 7 dpa as compared to the control. A vast number of DEGs, which are associated with proliferation and cell cycle, including *pcna*, *cdk2*, *cyclin D1 (ccnd1), lysineK-specific demethylase 8 (kdm8),* and *kinetochore protein* encoding *zwilch* were Up at 1 dpa and Down at 7 dpa, in line with the high proliferative activity at the early stage of melanocyte regeneration. Interestingly, most of the immune response-related genes including Fas receptor gene *fas*, *fyn-related Src family tyrosine kinase (frk), tumor necrosis factor b (tnfb),* and *complement component 4* (*c4*) appeared to be oppositely regulated at 1 dpa and 7 dpa (Figure 3A). Several neural crest-related genes including *paired box 3a (pax3a), SRY-box transcription factor 8b (Sox8b),* and *SRY-box transcription factor 10 (sox10) T*, which are also known for their role in pigment cell development (White and Zon 2008, Mort, Jackson et al. 2015), were Down at 1 dpa but Up at 7 dpa (Figure 3A). On the other hand, *regulator of G protein signaling 2 (rgs2)* and the transcription factor-encoding gene *forkhead box D3 (foxd3)*, act as negative regulators of neural crest development and melanocyte lineage development, respectively (Curran, Lister et al. 2010, Lin, Chiang et al. 2017), were Down at both 1 dpa and 7 dpa as compared to the control. *Rsg2* has been shown to restrict the *sox10(+)* non-ectomesenchymal lineage that acts as a source of melanocytes (Lin, Chiang et al. 2017). Thus, selective downregulation of *rsg2* and upregulation of *sox10* at 7 dpa support the idea that melanocyte lineage regeneration is promoted at the expense of the *sox10(-)* ectomesenchymal NCC. A large number of pigmentation-related genes such as *tyr*, *dct*, *oculocutaneous albinism II (oca2)*, *premelanosome protein a (pmela), endothelin 3b (edn3b),* and *tyrosinase-related protein 1a (tyrp1a)* (Cichorek, Wachulska et al. 2013) exhibited robust upregulation at 7 dpa, indicating induction of the melanocyte differentiation program (Figure 3A).

**Figure 3.**
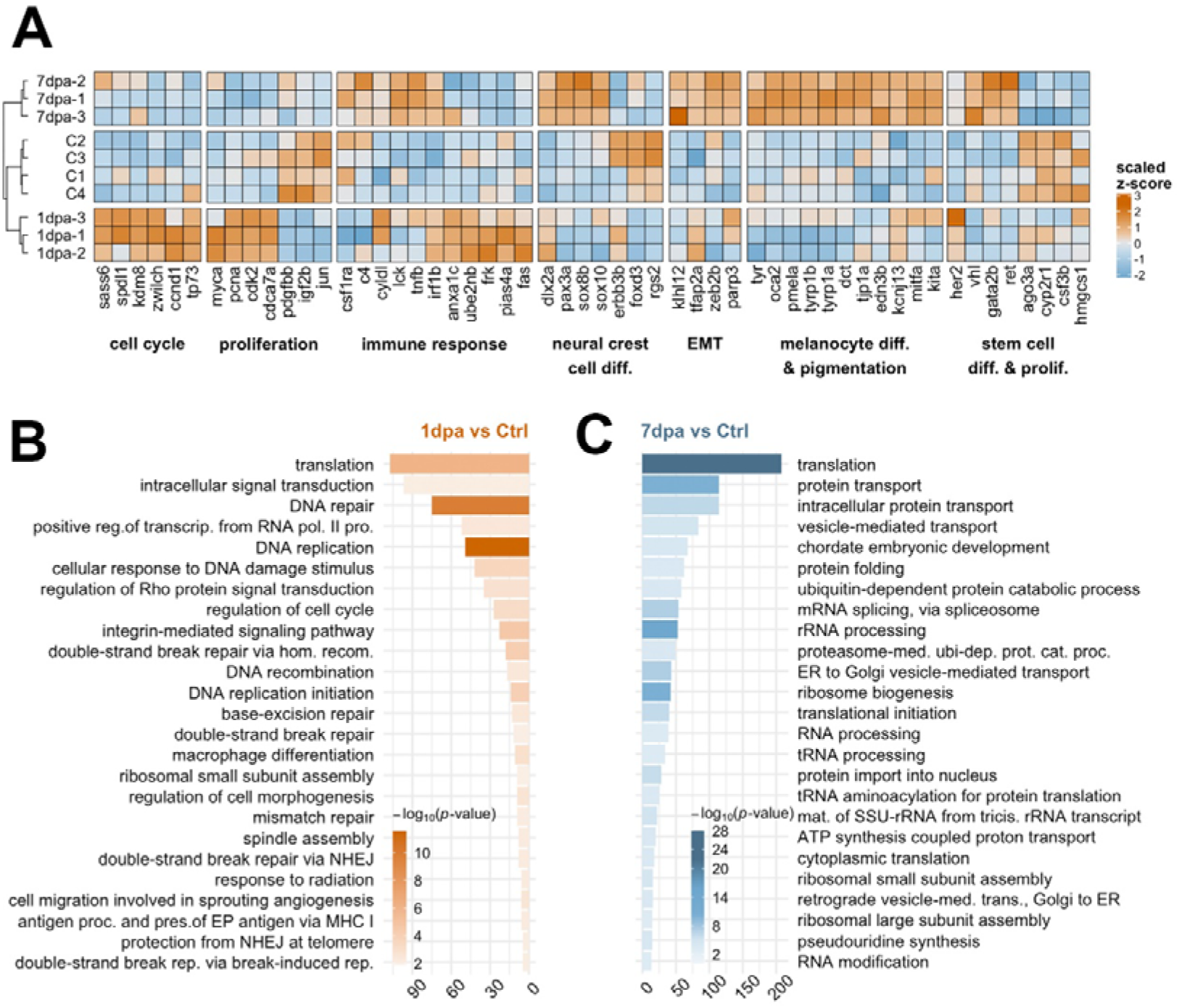
Early/proliferative and late/differentiation stages of melanocyte regeneration have distinct transcriptional profiles. **(A)** The heatmap shows the relative expression of selected genes between proliferation (1 dpa) and differentiation (7 dpa) stages together with the control group. Each column represents a single gene and each row represents a biological replicate of each condition. The scale bar shows z-scores of variance-stabilized (vst-transformed) counts from high to low expression, represented by the color gradient from orange to blue, respectively. **(B-C)** DAVID was used to show the 25 most significantly enriched GO-BP terms based on the transcriptional changes in 1 dpa and 7 dpa comparisons. The color scale shows -log_10_ of the EASE p-value and the x-axis shows the number of genes in each GO-BP term. dpa: days post-ablation, DAVID: Database for Annotation, Visualization, and Integrated Discovery, GO: Gene ontology, BP: Biological process.

Next, to further characterize the molecular signatures that define the early and late stages of melanocyte regeneration, we performed GO term enrichment analysis for 1 dpa and 7 dpa. At 1 dpa, the most significantly enriched DEGs were grouped in the biological process (BP) terms including DNA replication, DNA damage response, and various DNA repair mechanisms (Figure 3B, Table S3). KEGG pathway analysis and GSEA likewise revealed that, at 1 dpa, DEGs were positively enriched in DNA replication, cell cycle, and major DNA repair pathways (Figure S3A-B, Table S4). On the other hand, at 7 dpa, the DEGs were mostly enriched in GO-BP terms related to translation and RNA processing such as protein transport, protein folding, mRNA splicing, and rRNA processing (Figure 3C, Table S3). Top KEGG pathways and GSEA enriched at 7 dpa included protein translation, protein transport, general RNA processing, metabolic pathways, proteasome, and cellular respiration, mostly overlapping with the GO terms enriched at this stage (Figures S3A, S3C; Table S4). We also validated changes in the expression of the DEGs that are regulated at either 1 dpa or 7 dpa or both stages by qPCR (Figure S3D, Table S1). Thus, the early and late stages of melanocyte regeneration are considerably different from each other with respect to their transcriptome profiles.

### Transcriptional profile of melanoma differs from that of nevi by the proliferative burst and oppositely regulated NCC signature

To generate the melanoma model, we exploited the Tg*(mitfa:Hsa.HRAS^G12V^,mitfa:GFP)* zebrafish. In this line, the *mitfa* promoter drives the mutant *RAS* oncogene expression in the melanocyte lineage to promote the malignant transformation of melanocytes to melanoma, without the need for additional inactivating mutations on tumor suppressor genes (Santoriello, Gennaro et al. 2010, Forbes, Beare et al. 2017). As we could validate the zebrafish caudal fin as a reliable model for melanocyte regeneration, we decided to include the melanoma tissue that developed only on the caudal fin. In this way, we aimed to restrict the source tissue for transcriptome analysis to the caudal fin in both regeneration and cancer and minimize potential variation that might occur across different tissues. Thus, we collected tissues of nevi and melanoma that developed in the caudal fin of Tg*(mitfa+/-; mitfa:Hsa.HRAS^G12V^,mitfa:GFP)* (abbr. Tg *(mitfa+/-,HRAS^G12V^:GFP))* zebrafish for RNA-seq analysis (Figure 4A). PCA exhibited a very clear distinction between the control, nevi, and melanoma samples (Figure 4B). While only 1744 genes (1178 Up and 566 Down) were differentially expressed in nevi, the melanoma tissue harbored 11. 265 DEGs (5638 Up and 5627 Down) as compared to the control fin tissue (Figure 4C-D, Table S2).

**Figure 4.**
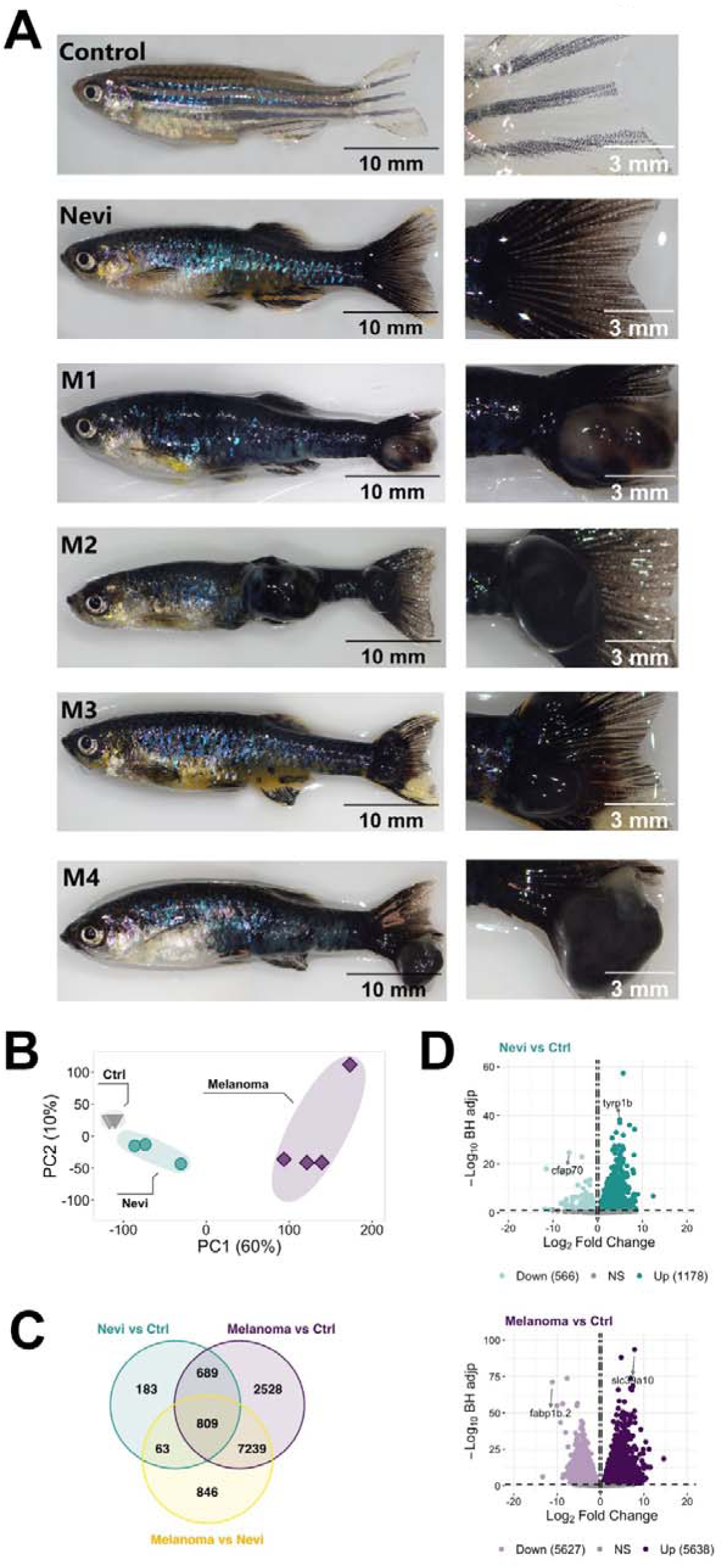
Validation of the melanoma model and quantitative analysis of the DEGs at the nevi and melanoma stages. **(A)** Zebrafish nevi and melanoma models. The whole fish are shown on the left and their caudal fins that were used for collection of the nevi and melanoma tissue are shown on the right. For melanoma samples, nodular melanoma samples were selected. Tissue materials were collected from the caudal fins of four individuals for each group and used as biological replicates (no pooling). Scale bars are 10 mm (whole fish on the left) and 3 mm (close-ups on the right). **(B)** Principal component analysis (PCA) of nevi, melanoma, and their controls. Different colors of dots and triangles represent the sample condition. Three sample groups were well clustered among their replicates and well separated from other sample groups. Principal Component 1 (PC1, x-axis) represents 60% and PC2 (y-axis) represents 10% of the total variation in the data. **(C)** The Venn diagram shows the number of differentially expressed genes (DEGs) in nevi vs control (turquoise), melanoma vs control (purple), melanoma vs nevi (yellow), and the overlap between each set of DEGs. **(D)** Volcano plots represent changes in the gene expression levels in nevi and melanoma. Each circle represents a gene. The x-axis shows log_2_ of fold change in the condition compared to the control and the vertical dashed lines indicate the fold change cutoffs. The y-axis depicts -log_10_ of the Benjamini-Hochberg adjusted p-value, where the significance threshold is indicated by a dashed horizontal line. Darker and lighter shades indicate upregulation and downregulation, respectively.

To further distinguish malignant melanoma from melanocytic nevi at the transcriptional level, we generated a heatmap of selected genes from among the most significant DEGs involved in proliferation/cell cycle, NCC differentiation, and melanocyte differentiation/pigmentation (Figure 5A). The control and nevi samples were similar in the expression of the majority of the selected genes that belong to proliferation and NCC differentiation groups. On the other hand, proliferation-related genes including *cdk2, minichromosome maintenance complex component 7 (mcm7), pcna, marker of proliferation Ki-67 (mki67),* and *cdk1* were Up in the melanoma tissues while being Down in nevi. The expression of the selected NCC gene signature was likewise completely oppositely regulated between control and melanoma samples. For example, neural crest-related genes including *erbb2, erbb3a, erbb3b, sox9b, twist1a, twist1b, and twist3* were strongly Down in melanoma while remaining Up in nevi as in control. These results suggest that the NCC signature has a potentially key role in the regulation of the malignant transformation of melanocytes to melanoma. For the melanocyte differentiation genes, we observed a similar pattern of regulation for nevi and melanoma. GO term enrichment analysis displayed that nevi were mostly enriched for GO-BP terms including various types of tissue development, pigmentation, and differentiation (Figure 5B, Table S3). KEGG pathway analysis for nevi showed enrichment of several metabolic pathways as well as melanogenesis, which regulates melanin production in a specialized organelle called melanosome (Figure S4A, Table S4). GSEA revealed that various pathways related to metabolism and melanogenesis were Up, while several signaling pathways including Wnt and TGF-β were Down in nevi samples (Figure S4B). Melanoma samples were significantly enriched for the GO-BP terms related to cell proliferation, cell migration, neural crest cell migration, development, and regeneration (Figure 5C, Table S3), and for the KEGG pathways that harbored numerous cells metabolism activities and cancer-associated signaling pathways such as MAPK, p53, FoxO and ErbB (Figure S4A, Table S4). According to GSEA, proliferation, and metabolic pathways were Up and several other signaling pathways including Wnt, TGF-β, Notch, GnRH, and calcium were Down in melanoma samples (Figure S4C). qPCR results confirmed differential expression of the genes that are regulated at either nevi or melanoma or both samples (Figure S4D, Table S1). Collectively, these data indicate that nevi and melanoma are similar concerning the induction of melanocyte differentiation and pigmentation responses and differ from each other regarding proliferation and NCC differentiation.

**Figure 5.**
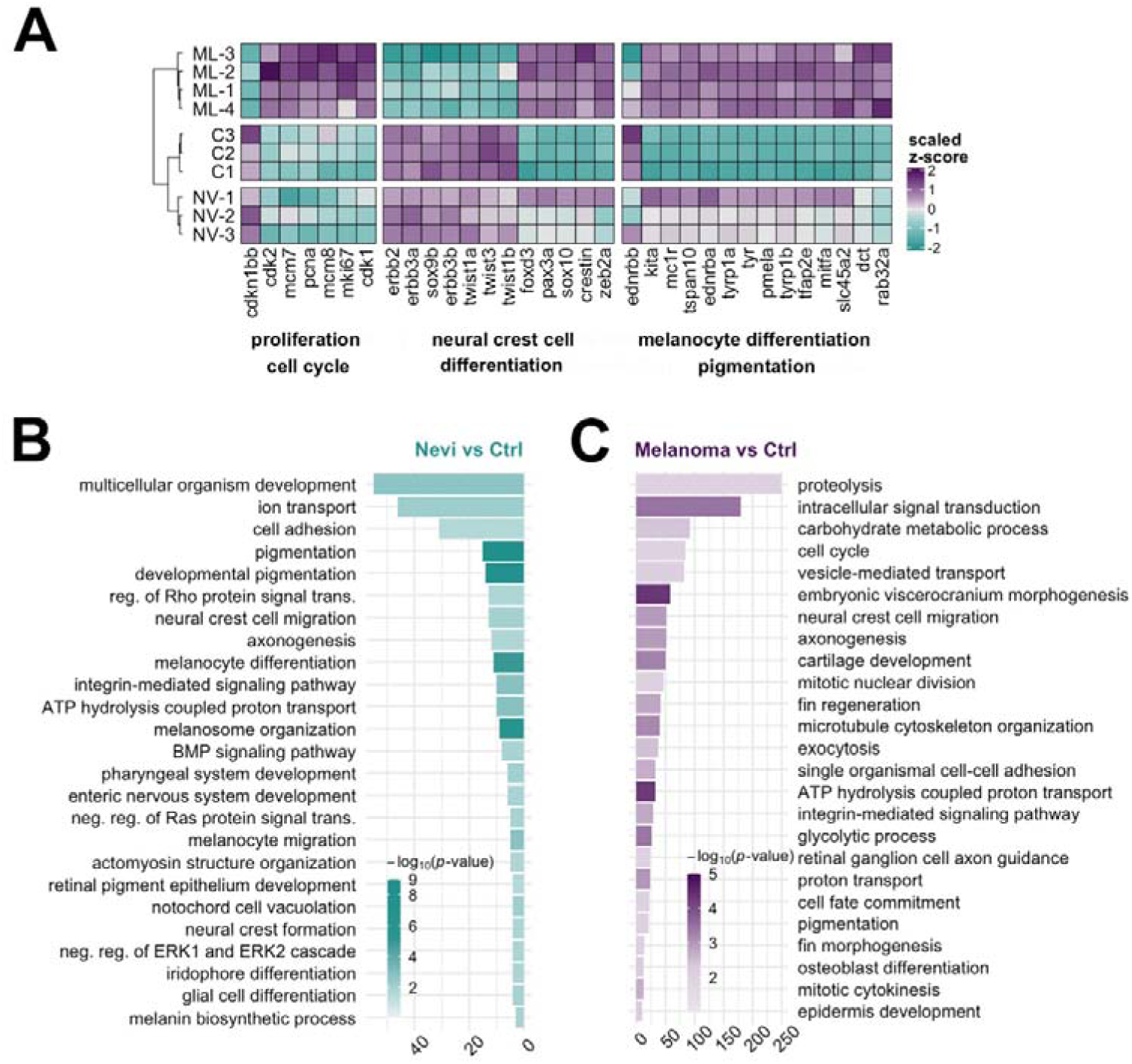
Transcriptional profile of melanoma differs from that of nevi by the proliferative burst and oppositely regulated NCC signature. **(A)** The heatmap shows the relative expression of selected genes between nevi and melanoma stages together with the control group. Each column represents a single gene, and each row represents a biological replicate of each condition. The scale bar shows z-scores of vst-transformed counts from high to low expression, represented by a color gradient from purple to turquoise, respectively. **(B-C)** DAVID was used to show the 25 most significantly enriched GO-BP terms based on the transcriptional changes in nevi and melanoma comparisons. The color scale shows -log_10_ of the EASE p-value and the x-axis shows the number of genes in each GO-BP term. DAVID: Database for Annotation, Visualization, and Integrated Discovery, GO: Gene ontology, BP: Biological process.

### Cellular processes including Wnt and TGF-***β***/BMP signaling pathways are differentially regulated between melanocyte regeneration and melanoma

As the cellular mechanisms of regeneration and cancer display shared and distinct patterns, next, we set out to compare the transcriptomes of the regenerating melanocytes to those of the nevi and melanoma. At this point, a global comparison would reveal how comparable regeneration and cancer are. Thus, by exploiting the GOChord function, we plotted a circularly composited overview of the fold changes of genes in the selected GO terms. To compare the changes in gene expression associated with these selected GO terms in the regeneration and cancer samples, we intersected the genes annotated in these terms with the DEG sets (Figure 6A). The genes obtained from the GO terms included a substantial number of genes involved in NCC migration, glial cell differentiation, fin regeneration, and fin morphogenesis. Interestingly, there were groups of DEGs that are regulated oppositely between 7 dpa and melanoma and other groups of DEGs that are regulated oppositely between the two regeneration stages (1 dpa and 7 dpa) and melanoma. Similar groups of genes were present in the GOChord plots generated by using the selected KEGG pathways (Figure S5). Moreover, we unraveled the DEGs that are shared between regeneration and cancer (1 dpa-nevi, 7 dpa-nevi, 1dpa-melanoma, and 7 dpa-melanoma) according to four different ways of regulation, i.e. Down-Up, Up-Up, Down-Down, Up-Down (Figure S6).

**Figure 6.**
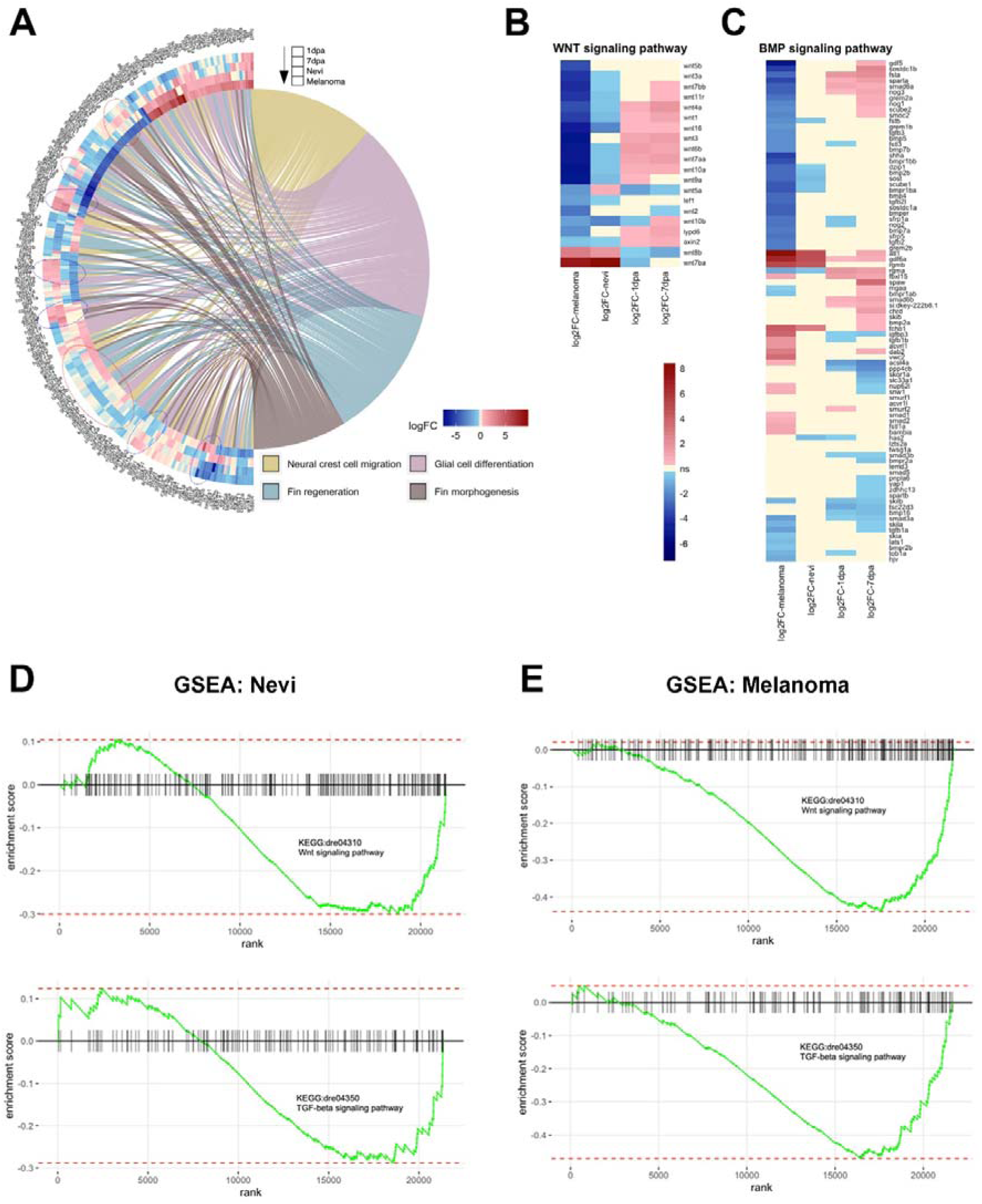
Cellular processes including Wnt and TGF-β/BMP signaling pathways are differentially regulated between melanocyte regeneration and melanoma. **(A)** GOChord plot shows log_2_ fold changes of the genes annotated in selected GO-BP terms for two stages of the melanocyte regeneration, nevi, and melanoma. The genes are linked to their assigned pathways by ribbons and ordered according to their log_2_ fold change values from low to high regulation, represented by a color gradient from blue to red, respectively. log_2_ fold changes are shown from the outer to the inner annulus in the following order: 1 dpa, 7 dpa, nevi, and melanoma. Blue dotted circles show clusters of DEGs that are oppositely regulated between 1 dpa/7 dpa and nevi/melanoma. Red dotted circles show clusters of DEGs that are oppositely regulated between 7 dpa and nevi/melanoma. **(B-C)** Heatmaps of selected genes in Wnt and BMP signaling pathways. Genes and conditions are clustered by the similarity of their differential expression profiles (log_2_ fold change). log_2_ fold change values are represented from high to low regulation by a color gradient from red to blue, respectively. Beige color represents non-significant differential expression (FC<1.2 in both directions or FDR>0.1). (D) GSEA plots of Wnt and TGF-β signaling pathways for zebrafish nevi samples. (E) GSEA plots of Wnt and TGF-β signaling pathways for zebrafish melanoma samples. GSEA was performed and enrichment plots were generated for selected gene sets using the fgsea package. On each enrichment plot, the horizontal black line represents the p-value ranks of genes with the most significant p-value rank on the left. The vertical black bars represent individual genes in the gene set and their ranks. The green curves represent the cumulative enrichment score (ES), and the red horizontal dashed lines show minimal and maximal scores. dpa: days post-ablation.

Next, we generated heatmaps of the DEGs that belong to two particular GO terms, i.e. Wnt signaling pathway and BMP signaling pathway. The majority of the Wnt pathway-related genes including *wnt7bb*, *wnt11r*, *wnt4a*, *wnt1*, *wnt16*, *wnt3*, *wnt6b*, *wnt7aa*, *wnt10a*, *lypd6,* and *axin2* were significantly Up in 1 dpa and 7 dpa, but Down in nevi and melanoma (Figure 6B). More than half of the genes that are associated with the BMP signaling pathway including *gdf5*, *smad6a*, *grem2a*, *BMP5*, *BMP7b*, *BMP2b*, *sost*, *BMP4*, *tgfb2l*, *BMP7a,* and *rgma* were likewise Down in melanoma, while not being significantly regulated or Up in 1 dpa and 7 dpa (Figure 6C). GSEA plots further validated that genes belonging to Wnt and TGF-β signaling pathways are among the strongly differentially expressed genes in nevi and, even more prominently, in melanoma (Figure 6D-E). TGF-β superfamily is a large group of structurally similar proteins and encompasses BMPs along with other growth and differentiation factors (Morikawa, Derynck et al. 2016). TGF-β and BMP signaling pathways, initiated with the binding of TGFβ/BMP ligands to type I and II receptors, can interact with each other at multiple levels including ligands, receptors, and downstream signaling components (Guo and Wang 2009, Dituri, Cossu et al. 2019). For instance, although BMPs predominantly signal through Smad1/5/8, some studies have reported BMP-induced activation of Smad2/3, which is typically associated with the TGF-β signaling pathway (Foletta, Lim et al. 2003, Holtzhausen, Golzio et al. 2014, Budi, Duan et al. 2017, Wnuk, Paw et al. 2020). Moreover, TGF-β and BMP signaling pathways have been shown to activate each other in some cellular contexts (Chen, Deng et al. 2012). Thus, these results showing enrichment of genes related to both TGF-β and BMP signaling pathways propose crosstalk of these pathways also in melanoma development. Overall, our data strongly suggest that Wnt and TGF-β/BMP signaling pathways are differentially regulated between melanocyte regeneration and melanoma.

### Activation of canonical Wnt or BMP signaling pathways enhances larval melanocyte regeneration

Because of the differential regulation of Wnt and TGF-β/BMP signaling pathways between melanocyte regeneration and melanoma, we used drugs that inhibit or activate these pathways and first assess the influence on the melanocytes during regeneration. Since most of the Wnt pathway-related genes, including *wnt7bb*, *wnt1*, *wnt3*, *lypd6,* and *axin2* that are regulated oppositely between melanocyte regeneration and melanoma, have been associated with the Wnt/β-catenin pathway, we decided to modify the β-catenin-dependent (canonical) Wnt signaling among the Wnt pathways (Jho, Zhang et al. 2002, Grumolato, Liu et al. 2010, Özhan, Sezgin et al. 2013, Arensman, Kovochich et al. 2014). We exploited the larval zebrafish that is rapidly responsive, easy to manipulate, and permeable to small molecules.

First, to assess the influence of canonical Wnt and TGF-β/BMP signaling pathways on melanocyte regeneration, we ablated the melanocytes of the zebrafish larvae with 4-HA from 36 hpf to 60 hpf and treated the larvae with the Wnt antagonist IWR, Wnt agonist BIO, BMP antagonist DMH1 or TGF-β/BMP agonist ISL from 60 hpf to 4 dpf (Figure 7A). Upon 4-HA treatment, melanocytes were lost as early as 48 hpf (Figure 7B). By 4 dpf, 4-HA significantly reduced the number of melanocytes, which further decreased after the inhibition of either canonical Wnt or BMP pathways with IWR or DMH1, respectively (Figure S7A-D). In contrast, the activation of either pathway with BIO or ISL increased the number of melanocytes in 4-HA-treated larvae (Figure 7C-D). As 4-HA-mediated melanocyte ablation does not affect the MSCs in zebrafish, the increase in the number of melanocytes is most likely due to the melanocyte regeneration promoted by the activation of either pathway (O’Reilly-Pol and Johnson 2013). There was no detectable alteration in the number of newly produced melanocytes in the larvae treated with activators of canonical Wnt and TGF-β/BMP pathways (Figure S7D-E), while both pathways were activated as revealed by elevation of phospho-β-catenin (phosphorylated at Ser675, increasing nuclear localization and transcriptional activation of β-catenin) or TGF-β/BMP target genes *runx2* and *sp7 (osterix)* (Lee, Kim et al. 2000, Zhang, Zhang et al. 2019) (Figure S7F-G). Thus, activation of canonical Wnt and TGF-β/BMP pathways can promote the regenerative capacity of the melanocytes.

**Figure 7.**
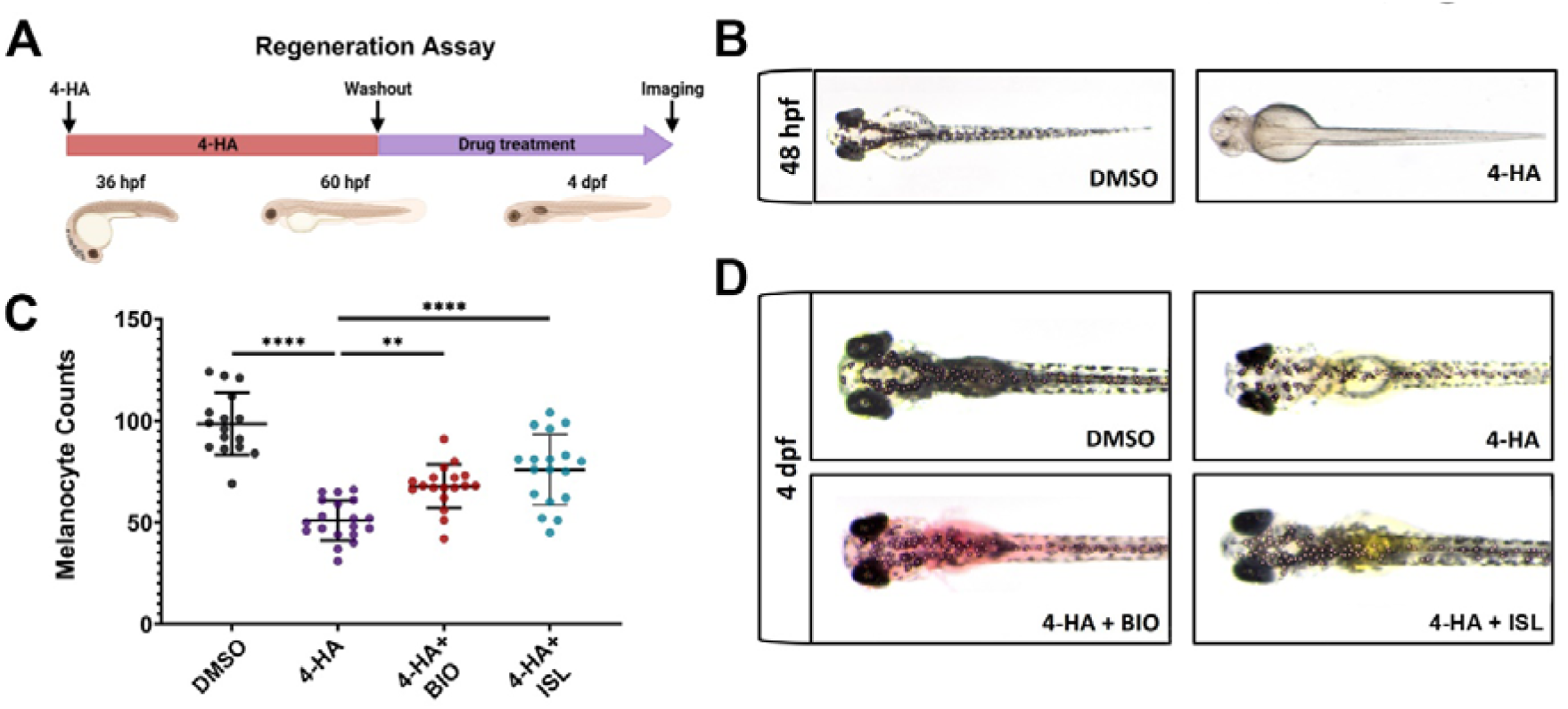
Activation of canonical Wnt or TGF-β/BMP signaling pathways enhances larval melanocyte regeneration. **(A)** Scheme for experimental design of melanocyte regeneration. Zebrafish embryos were treated with 4-HA from 36 hpf to 60 hpf. After washout of 4-HA at 60 hpf, drugs were administered into the embryo water, and larvae were analyzed at 4 dpf. **(B)** 2 dpf zebrafish larvae treated with DMSO or 4-HA. **(C)** Dot plot showing the number of melanocytes in 4 dpf larvae treated with DMSO, 4-HA, 4-HA+BIO, or 4-HA+ISL. Each dot represents one larva (DMSO n=16, 4-HA n=19, 4-HA+BIO n=18, 4-HA+ISL n=17). Statistical significance was evaluated using a one-way ANOVA test **p<0.01 and ****p < 0.0001. **(D)** Representative images of melanocyte regeneration groups are counted in (C).

### Activation of canonical Wnt or TGF-***β***/BMP signaling pathways suppresses invasiveness, migration, and proliferation of human melanoma cells

As canonical Wnt or TGF-β/BMP signaling pathways were downregulated in melanoma and, when activated, enhanced melanocyte regeneration, we next aimed to test whether activation of either pathway exerts anti-cancer effects on the SK-MEL-28, a human malignant melanoma cell line. Initially, to investigate whether EMT was affected in these invasive tumor cells upon BIO or ISL treatment, we examined actin stress fiber formation using phalloidin staining (Figure 8A). We observed that DMSO-treated melanoma cells had actin stress fibers that were aligned and bundled along their length, consistent with a mesenchymal phenotype (Figure 8A). BIO or ISL treatment, however, resulted in disorganization and loss of this aligned shape of stress fibers, a decrease in their overall length, and their disorganization. There was a concomitant decrease in the mRNA expression levels of the EMT marker genes *N*-*cadherin*, *Snail, Slug,* and *Zeb1* (Figure 8B). Moreover, vimentin, a major component of mesenchymal cells, was cleaved in drug-treated melanoma cells, suggesting disruption of their cytoskeletal structure and inhibition of their migration (Figure 8C). Finally, scratch wound healing assay revealed that activation of Wnt/β-catenin or TGF-β/BMP signaling caused significant retardation of gap closure rate, further supporting the inhibitory influence of both signaling pathways on cancer cell migration (Figure 8D-E).

**Figure 8.**
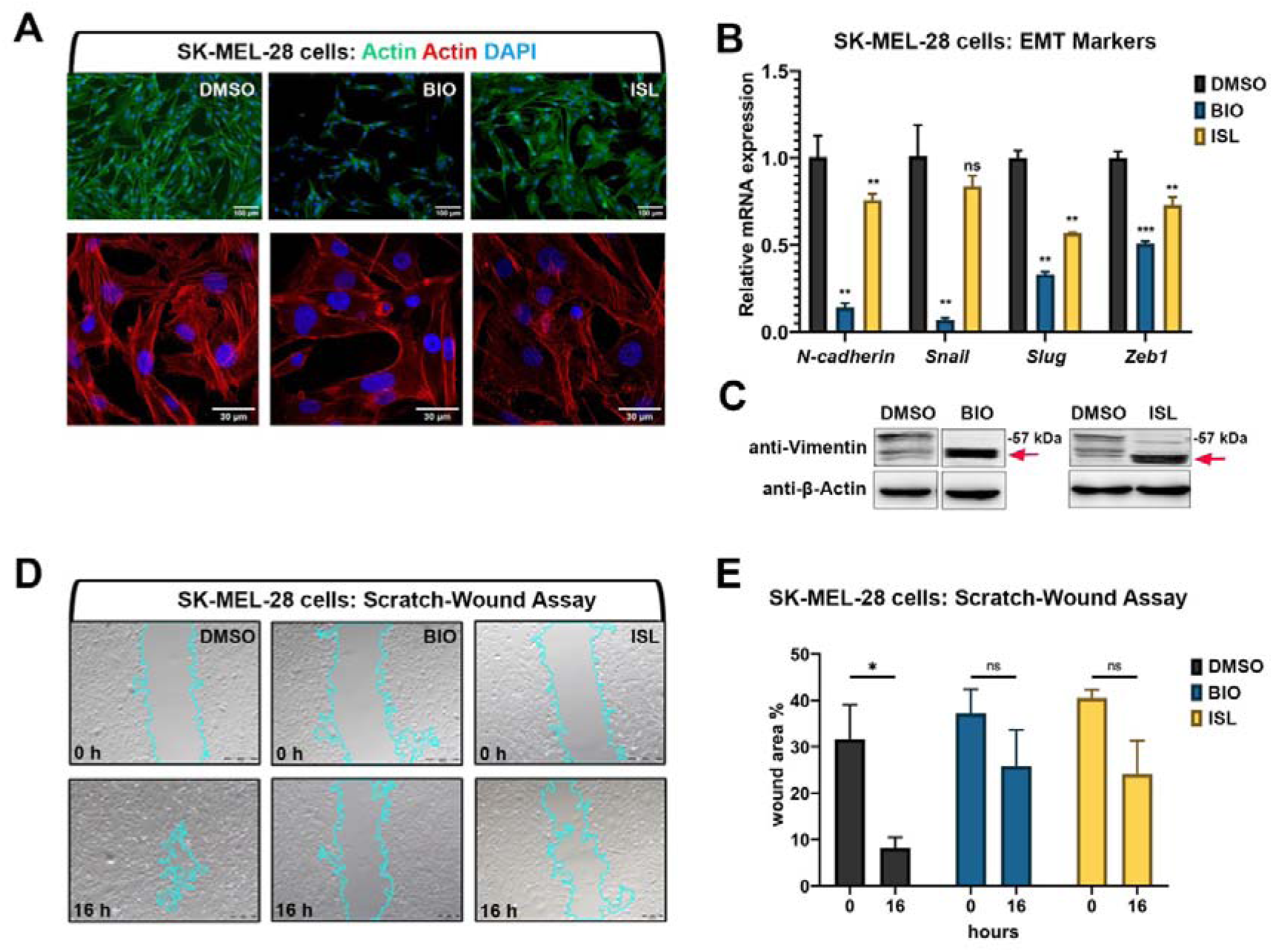
Activation of canonical Wnt or TGF-β/BMP signaling pathways suppresses invasiveness of human melanoma cells. **(A)** Phalloidin staining in SK-MEL-28 melanoma cell line treated with BIO or ISL reveals changes in actin stress fiber formation. **(B)** qPCR on SKMEL-28 cells treated with BIO or ISL showing expression of EMT marker genes *N-cadherin*, *Snail*, *Slug,* and *Zeb1*. *GAPDH* was used as the housekeeping control gene. Error bars represent ± standard error of the mean (SEM, n=3). Statistical significance was evaluated using an unpaired t-test. **p<0.01, ***p<0.001 and ns: nonsignificant. **(C)** Western blot of SKMEL-28 cells treated with BIO or ISL for the mesenchymal marker vimentin. Red arrow indicates cleaved vimentin. **(D)** Scratch-wound assay in SKMEL-28 cells treated with BIO or ISL. **(E)** Wound closure expressed as the percentage of the remaining area not covered by the cells 16 hours after the scratch. Error bars represent ± standard error of the mean (SEM, n=3). Statistical significance was evaluated using an unpaired t-test. *p<0.05, **p<0.01, ***p<0.001 and ns: non-significant.

Next, to examine the effect of Wnt/β-catenin and TGF-β/BMP pathways on melanoma concerning cancer cell migration and metastasis, we exploited larval zebrafish xenografts by injecting DiO-labeled SK-MEL-28 cells into the yolk sac at 2 dpf, treated them with the agonists of the pathways and observed the cancer cell behavior over time. At 5 dpi, melanoma cells invaded the caudal tissue and formed micrometastasis colonies in DMSO-treated control embryos (Figure 9A-B). On the other hand, activation of the canonical Wnt (with BIO) or TGF-β/BMP (with ISL) pathway significantly reduced their dissemination through the caudal tissue. To assess whether the reduction in the number of migrating cells results from an alteration in cell proliferation and/or cell death, we further evaluated cell proliferation and cell death by apoptosis in the larval xenografts. Whole-mount immunofluorescence and confocal imaging quantification of DiO+, DAPI+ SK-MEL-28 cells revealed that BIO or ISL-treated xenografts displayed dramatically lower rates of melanoma cell proliferation, indicated by the percentage of mitotic figures (Figure 9C-D). On the other hand, apoptosis of DiO+, DAPI+ SK-MEL-28 cells did not alter significantly in response to any drug treatment (Figure S8A-B). To further dissect the impact of the activation of canonical Wnt or TGF-β/BMP signaling on melanoma, we took advantage of the Tg*(mitfa:Hsa.HRAS^G12V^,mitfa:GFP)* line, which was outcrossed to wt AB zebrafish. As 50% of the offspring had the Tg *(mitfa+/-,HRAS^G12V^:GFP)* genotype and their growing melanocytes specifically expressed *RAS^G12V^* oncogene, the number of melanocytes dramatically increased in these larvae within 5 days after fertilization, resulting in melanocytic nevi formation. Activation of either signaling pathway with BIO or ISL efficiently reduced the levels of phosphorylated ERK (p-ERK), a major downstream readout of activated Ras/MAPK signaling in Tg*(mitfa+/-,HRAS^G12V^:GFP)* larvae at 5 dpf (Figure 9E). In addition, BIO or ISL treatment reduced the expression of late melanocyte differentiation markers *tyr* and *dct* in Tg*(mitfa+/-,HRAS^G12V^:GFP)* larvae as compared to Tg(*mitfa+/-*) control larvae at 5 dpf (Figure 9F). Finally, the melanin content assay confirmed that the number of melanocytes was significantly reduced in Tg*(mitfa+/-,HRAS^G12V^:GFP)* larvae when compared to the control (Figure 9G). These results strongly suggest that activation of canonical Wnt or TGF-β/BMP signaling can block endogenous melanoma growth. Together, these data suggest that activation of canonical Wnt and TGF-β/BMP pathways can efficiently suppress the invasive, migratory, and proliferative potential of human melanoma cells *in vitro* and *in vivo*.

**Figure 9.**
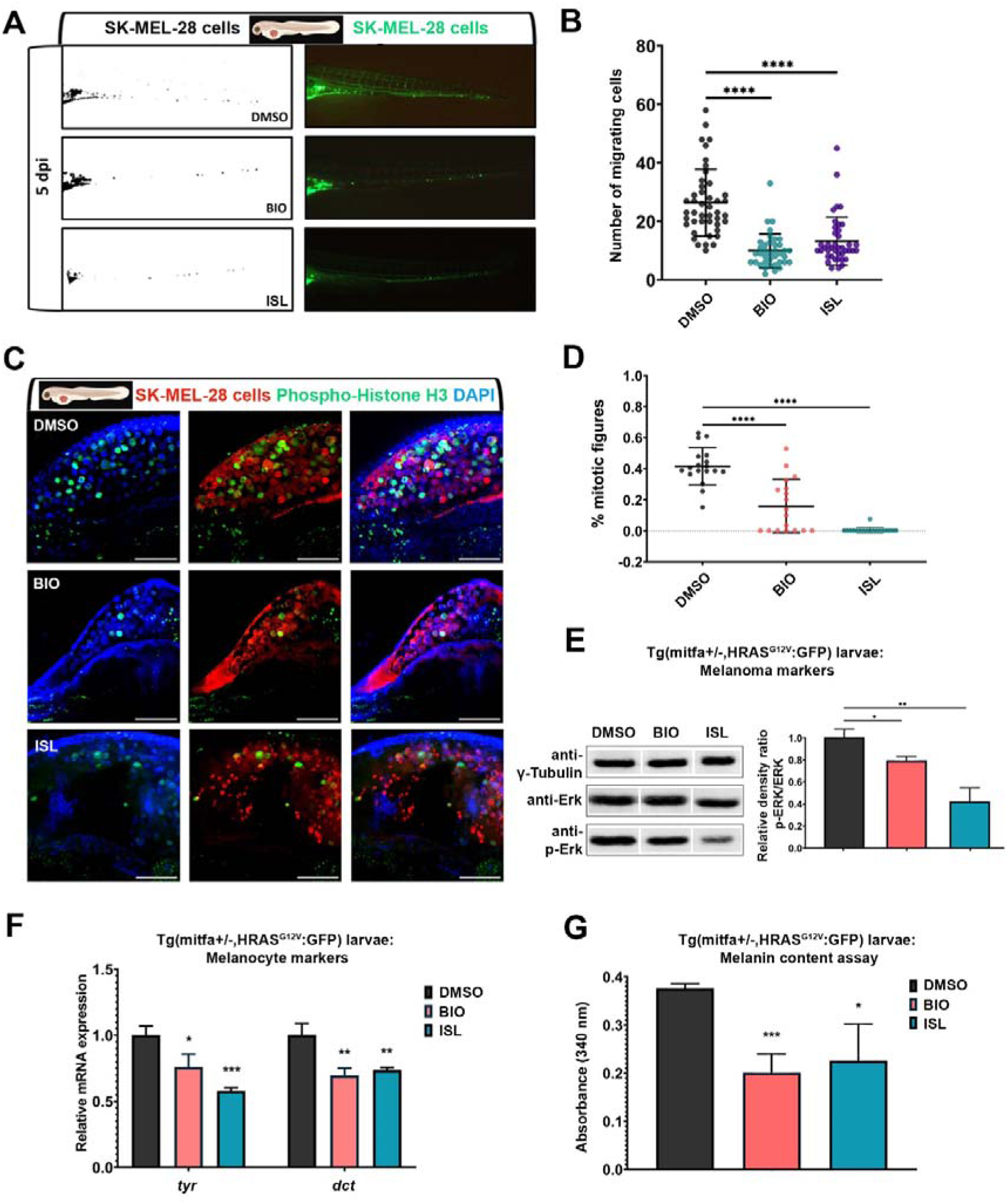
Activation of canonical Wnt or TGF-β/BMP signaling pathways suppresses migration and proliferation of human melanoma cells *in vivo*. **(A)** Representative fluorescence microscope images of 7 dpf zebrafish larvae xenografted with human melanoma cells (SK-MEL-28) at 2 dpf and treated with DMSO, BIO or ISL. **(B)** Dot plot showing the number of invading cells at 7 dpf larval xenografts with micrometastasis. Each dot represents one larval xenograft (DMSO n=44, BIO n=38, ISL n=41). **(C)** Representative confocal microscope images of anti-phospho-histone H3 (green) staining of 7 dpf zebrafish larvae xenografted with SK-MEL-28 cells (red) at 2 dpf and treated with DMSO, BIO, or ISL. **(D)** Dot plot showing the percentage of mitotic figures in each treatment. Each dot represents mitotic figures counted in each z-stack slice divided by the number of DiO+, DAPI+ nuclei in (C). Larvae were counterstained for DAPI. Scale bars 50μm. Statistical significance in B and D was evaluated using a one-way ANOVA test ****p < 0.0001. **(E)** Western blot of 5 dpf Tg*(mitfa+/-,HRAS^G12V^:GFP)* zebrafish larvae treated with BIO or ISL for total and phospho-ERK (p-ERK). Tg(*mitfa+/-*) larvae were used as control. Graph shows the average relative density ratios of p-ERK to total ERK from three independent experiments. **(F)** qPCR on Tg*(mitfa+/-,HRAS^G12V^:GFP)* zebrafish larvae treated with BIO or ISL showing reduced expression of pigmentation marker genes *tyr* and *dct* at 5 dpf. Tg(*mitfa+/-*) larvae were used as control. *rpl13* was used as the housekeeping control gene. **(G)** Melanin content of Tg*(mitfa+/-,HRAS^G12V^:GFP)* zebrafish larvae treated with BIO or ISL. Error bars represent ± standard error of the mean (SEM, n=6). Statistical significance was evaluated using an unpaired t-test. *p_≤_0.05, **p_≤_0.01 and ***p_≤_0.001.

## Discussion

Tissue regeneration mechanisms have been discussed from the perspective of differential gene expression analysis in different tissues. However, there is limited information on the comparative analysis of the transcriptomes in different stages of regeneration and the dynamic transcriptional responses during regeneration. Recently, we have revealed the transcriptomes of the regenerating zebrafish brain at its two different stages (Demirci, Cucun et al. 2020). Here, we first extended this approach to reveal the transcriptome profiles of zebrafish melanocyte regeneration at two different stages, i.e. at 1 dpa and 7 dpa that represent the early/proliferative and the late/differentiation stages, respectively. Since the molecular pathways and gene expression patterns of regeneration show similarities to those of carcinogenesis, a global comparison of the transcriptomes during both phenomena could provide a large amount of information on the common and distinct mechanisms of regeneration and cancer. Thus, next, by implementing this viewpoint to the melanocyte model, we generated zebrafish models of early and late melanoma and compared their transcriptomes to each other and those of the regenerating melanocytes. We used the melanocytic nevi to represent the early stage of melanoma since a considerable percentage of all melanoma cases are associated with preexisting dysplastic moles (nevi), where MAPK-related genetic alterations are responsible for hyper-proliferation of the skin melanocytes (Bevona, Goggins et al. 2003). Thus, Tg*(mitfa:Hsa.HRAS_G12V,mitfa:GFP)* zebrafish line that develops melanocytic nevi is a valuable model to investigate melanoma transformation from the earliest steps. Considering the importance of cell-cell and cell-extracellular matrix interactions in the architecture of both stem/progenitor cell niches during regeneration and tumor formation, we aimed to obtain a broad overview of the average trends in gene expression profiles of cell populations involved in regeneration and cancer. Thus, we exploited bulk tissue RNA-seq to identify the differences between sample conditions (i.e. early-regenerating, late-regenerating, early cancer, and late cancer) by measuring the average global gene expression differences across the population of cells. To our knowledge, this is the first study that compares the molecular mechanisms of regeneration and cancer that occur within the same tissue and the same organism. Accordingly, we have reached the following conclusions: i) 1 dpa represents the proliferation stage and is used to analyze the early regeneration, while 7 dpa represents the differentiation stage and is used to examine the late regeneration at the transcriptional level. These stages significantly differ from each other concerning the clustering and content of the DEGs. ii) While 1 dpa is enriched with DEGs related to proliferation and cell cycle, 7 dpa is mostly characterized by DEGs associated with melanocyte differentiation and pigmentation. Genes related to immune response and NCC differentiation were generally oppositely regulated between 1 dpa and 7 dpa. iii) Selected DEGs related to proliferation, cell cycle, and NCC differentiation are mostly oppositely regulated in samples of melanocytic nevi and malignant melanoma. On the other hand, these genes show similar expression patterns between nevi and control. Genes associated with melanocyte differentiation and pigmentation are significantly upregulated in nevi and even more in melanoma, potentially due to the higher number of melanocytic cells in the melanoma samples as compared to the nevi. iv) Genes belonging to the Wnt signaling pathway are mostly oppositely regulated between regeneration and cancer; i.e. majority of them are upregulated during melanocyte regeneration but downregulated in melanoma. The majority of TGF-β/BMP signaling pathway-related genes are either oppositely regulated between regeneration and cancer or have not changed in one phenomenon while up/downregulated in the other one. v) During regeneration, activation of Wnt or TGF-β/BMP pathways significantly increased the number of newly produced melanocytes. In contrast, in melanoma, the activation of either pathway suppressed the invasiveness of cells and decreased the number of migrating and proliferating cells. Thus, these pathways appear to operate in favor of melanocyte regeneration while interfering with melanoma growth.

The early stage of regeneration (1dpa) and melanoma is characterized by the upregulation of genes including *cdk2, mcm7, pcna, mki67,* and *cdk1* that are associated with proliferation and cell cycle. On the contrary, these genes were likewise regulated between the late stage of regeneration (7 dpa) and control as well as between nevi and control. These data suggest that cell proliferation is a common signature between cancer and early regeneration, but not late regeneration. In contrast, *cdkn1bb*, the ortholog of human *CDKN1B*, was downregulated both at 1 dpa and in melanoma. By inhibiting the activity of Cdk-cyclin complexes at the G1 phase, *CDKN1B* negatively regulates cell proliferation (Polyak, Lee et al. 1994, Sharma and Pledger 2016). Cancers carrying BRAF mutations have increased expression of cell cycle inhibitors including *CDKN2A*, *CDKN1A,* and *CDKN1B* (Chu, Hengst et al. 2008). However, these genes have been reported to be recurrently inactivated in pre-malignant nevi with BRAF mutations and contribute to melanoma pathogenesis (Gruber, Kastelan et al. 2008, González-Ruiz, González-Moles et al. 2020). Thus, parallel downregulation of *cdkn1bb* during early melanocyte regeneration and melanoma suggests that this tumor suppressor gene may act as a key regulator, among others, making early regeneration and cancer comparable concerning proliferation signatures.

During early melanocyte regeneration, we found that in addition to the cell cycle and proliferation genes, DNA repair signatures were significantly enriched most likely because actively proliferating cells are exposed to DNA damage more than any other tissue. Strikingly, even though early melanocyte regeneration and melanoma are similar concerning the enrichment of genes related to cell proliferation, they look completely different in the activation of DNA repair mechanisms. Whereas early regeneration was enriched for various types of DNA repair including base excision, mismatch, and double-strand break, melanoma showed no enrichment for DNA repair pathways. This is also supported by the enrichment of the KEGG pathway called Fanconi anemia which has been shown to promote homologous recombination upon DNA interstrand crosslinks by suppressing nucleotide-excision repair and is mostly deregulated in cancer predispositions (Sancar and Tang 1993, Michl, Zimmer et al. 2016, Fang, Wu et al. 2020, Zhang, Mou et al. 2021). Proper activation of DNA repair mechanisms is essential for homeostatic renewal and tissue regeneration following injury since defects in these mechanisms could interfere with the ability of adult stem cells to rebuild healthy tissues and increase the susceptibility to malignant transformations (Al Zouabi and Bardin 2020). Thus, while maintaining an exquisite balance between proliferation and differentiation, tissue regeneration carries the risk of malignant transformations once the cells escape from the rigid control of the cell cycle and DNA repair mechanisms, especially at the early stages of regeneration.

NCC signature including *foxd3*, *pax3a*, *sox10*, *crestin,* and *zeb2a* showed a significant increase in both nevi and melanoma. The transcriptome profile of nevi samples appears to resemble that of the superficial spreading melanoma, where the stem cell-related genes *erbb2* and *erbb3* and the mesenchymal signatures *sox9, twist1a, twist1b,* and *twist3* showed increased expression (Buac, Xu et al. 2009, Travnickova, Wojciechowska et al. 2019). The upregulation of *crestin* is in line with its instant re-expression at the onset of melanoma (Kaufman, Mosimann et al. 2016, McConnell, Mito et al. 2019). On the other hand, *foxd3* acts as a repressor of *mitfa* and inhibits melanocyte differentiation during NCC development (Curran, Lister et al. 2010). Therefore, elevated *foxd3* expression in our melanoma samples could be related to the increase in stemness/less differentiated phenotype.

One of the most prominent effects shared between late melanocyte regeneration and melanoma is the evident upregulation of genes involved in melanocyte differentiation and pigmentation. One of those genes, the *mitfa* gene that is essential for the development of melanocytes, has been reported to have contradictory roles in melanoma. For example, *MITF* amplification is found in 15-20% of human metastatic melanomas with poor prognosis (Ugurel, Houben et al. 2007). MITF has been shown to directly regulate a set of genes required for cell proliferation and DNA repair and thus inhibit cellular senescence (Strub, Giuliano et al. 2011). Interestingly, both high and low levels of MITF in melanomas have been linked with resistance to chemotherapeutic agents, increased invasiveness as well as melanoma relapse after targeted therapy against BRAF mutations and MEK activity (Eccles, He et al. 2013, Müller, Krijgsman et al. 2014, Van Allen, Wagle et al. 2014). These paradoxical functions of MITF have been explained by the “MITF rheostat” model, where its high-level, mid-level, low-level, and absence promote differentiation, proliferation, invasion, and cell death, respectively (Carreira, Goodall et al. 2006, Hoek and Goding 2010, Seberg, Van Otterloo et al. 2017). Thus, increased expression of *mitfa* in our melanoma samples is likely associated with survival and proliferation of tumor cells as the samples were at the same time significantly enriched for the cell cycle and proliferation-related signatures. The key role of human *MITF* in melanocyte differentiation occurs through direct transcriptional control of several genes including TYR, TYRP1, DCT, KIT, MC1R, PMEL17, and SLC45A2 as well as indirect ones such as EDNRB (Sato-Jin, Nishimura et al. 2008, Cheli, Ohanna et al. 2010). We have found an array of pigmentation genes including *kita*, *dct*, *tyrp1a*, *tyrp1b, ednrba, ednrbb, mc1r, pmela,* and *slc45a2* to be significantly upregulated during late melanocyte regeneration and melanoma, strongly suggesting that their expression is transcriptionally controlled by *mitfa*. At this point, it is important to mention that we found Tg *(mitfa+/-,HRAS^G12V^:GFP)* zebrafish to develop tissues of amelanotic melanoma, a subtype of melanoma that has little to no pigment, which expresses relatively lower levels of *mitfa* than the pigmented melanoma tissue (data not shown). The human amelanotic melanoma A375 cells were also reported to express a very low level of *MITF* and its target gene *TYR* (Beuret, Flori et al. 2007). Therefore, it is likely that the level of *mitfa* is key to adjusting the target gene expression and determining the subtype of melanoma.

Our data suggest that regenerating tissue and tumor mass diverge concerning the regulation of canonical Wnt and TGF-β/BMP signaling pathways, especially in the late stages of regeneration and cancer. As both pathways showed a general tendency to be upregulated in melanocyte regeneration and downregulation in melanoma, it would be interesting to test whether they are necessary for regeneration and able to interfere with the malignant behavior of the cancer cells. Our functional analysis in the zebrafish larvae indeed showed that the activation of canonical Wnt or TGF-β/BMP pathways was essential for efficient melanocyte regeneration. These results have further supported the findings obtained from the loss-of-function mouse models, which dissected the collaborative role of Wnt and BMP signaling pathways in triggering the commitment of proliferative MSCs to differentiation in a MITF-dependent manner (Infarinato, Stewart et al. 2020). On the other hand, the activation of canonical Wnt or TGF-β/BMP signaling with BIO or ISL, respectively, could inhibit the invasive, migratory, and proliferative behavior of human melanoma cells *in vitro* and in the zebrafish xenograft model. Activation of Wnt/β-catenin signaling via GSK3 inhibitors including BIO, LY2090314, Chir98014, and Chir99021, has been revealed to block the migratory and invasive behavior of melanoma cells by downregulating the expression of N-cadherin and reduces their proliferation *in vitro* and *in vivo* (John, Paraiso et al. 2012, Atkinson, Rank et al. 2015, Taylor, Rothstein et al. 2018). Intriguingly, the GSK-3β-dependent Wnt pathway activation appears to reduce invasion and proliferation of melanoma cells *in vitro* and melanoma growth in xenografts *in vivo* by suppressing the expression of Sox10, which significantly increased in our nevi and melanoma samples (Uka, Britschgi et al. 2020). ISL has likewise been shown to exert anti-cancer effects on melanoma by suppressing cell proliferation, migration, invasion, and metastasis or inducing apoptosis in various melanoma cell lines (Xiang, Chen et al. 2018, Wang, Yu et al. 2021, Xiang, Zeng et al. 2021, Wu and Wang 2023). Since a cancer-promoting role of BMP signaling has been reported in a *BRAF^V600E^*-initiated melanoma model of zebrafish, differential biological effects of RAS and BRAF oncogenic signaling in the regulation of key pathways and genes should be taken into consideration (Oikonomou, Koustas et al. 2014, Gramann, Frantz et al. 2021).

In conclusion, we took advantage of modeling melanocyte regeneration and melanoma in zebrafish that can efficiently regenerate and be induced to form cancers in zebrafish, enabling regeneration and cancer to be successfully modeled and compared within the same organism. Moreover, by restricting the tissue for sample collection to the same organ, i.e. the caudal fin, which harbors melanocytes to efficiently model melanocyte regeneration and melanoma, we aimed to minimize the differences that might stem from the source tissue. Our detailed transcriptome analyses have revealed the common and distinct genes and pathways between melanocyte regeneration and melanoma at their early and late stages. Characterization of those pathways that secure proper initiation and termination of the proliferation and differentiation states during regeneration could be particularly helpful in the development of anticancer strategies. Regulatory genes and signaling pathways that are differentially regulated between regeneration and cancer may constitute useful starting points to test their potential to stop tumor growth and even reverse cancer progression. Our findings also highlight the necessity of a contextual understanding of these genes and pathways, including canonical Wnt and TGF-β/BMP signaling pathways, in specific tumor types, to construe their involvement in cancer progression and explore their therapeutic potential.

## Data Availability

All datasets have been deposited in ArrayExpress under the link: https://www.ebi.ac.uk/arrayexpress/experiments/E-MTAB-11163/ with the accession number “E-MTAB-11163”.

## Acknowledgment

We would like to thank Dr. Adam Hurlstone for the zebrafish lines *mitfa*-/-(*nacre*) and Tg*(mitfa:Hsa.HRAS^G12V^,mitfa:GFP)*. We would like to thank Izmir Biomedicine and Genome Center Vivarium-Zebrafish Core Facility, Optical Imaging Core Facility, and Histopathology Core Facility for providing zebrafish care, microscope facility support, and histopathology service support, respectively. We also thank the Genomics Core Facility (GeneCore) of EMBL, Heidelberg.

## Ethics Statement

The animal study was reviewed and approved by the Animal Experiments Local Ethics Committee of Izmir Biomedicine and Genome Center (IBG-AELEC).

## Author Contributions

GO and EK designed the experiments. EK and DK performed molecular and cell biology experiments. YD and GH conducted the bioinformatics analyses. GO and EK wrote the manuscript. All authors contributed to the discussion.

