## Supplementary Figures for "Canonical Wnt and TGF-β/BMP signaling enhance melanocyte regeneration and suppress invasiveness, migration, and proliferation of melanoma cells"

**Katkat et al. Supplementary Figures and Figure Legends**

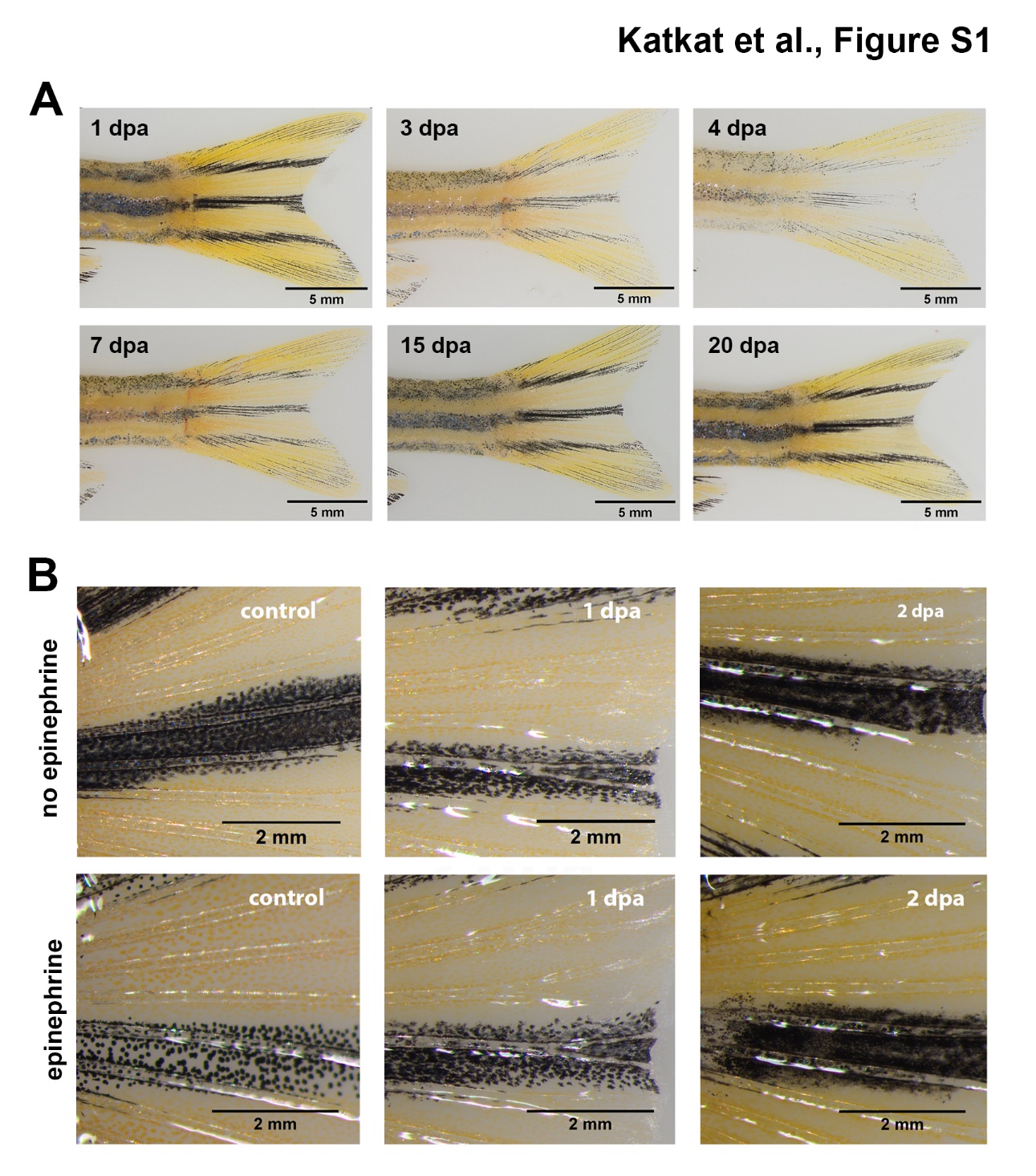

**Figure S1.** **Regeneration and epinephrine response of the zebrafish melanocytes (A)** Zebrafish melanocyte regeneration within 20 days after 1 day of NCP treatment. Loss of pigmentation was visible at 3 dpa. Melanocytes were totally disposed out of the skin at 4 dpa. Melanocyte re-pigmentation was detectable at 7 dpa. Scale bars are 5 mm. **(B)** Caudal fins of adult zebrafish following with or without epinephrine treatment. Live melanocytes of the control group rapidly responded to epinephrine by losing their dendritic shape and gathering melanosomes around the nucleus. NCP-treated fish showed no response to epinephrine at 1 dpa and 2 dpa, indicating that their melanocytes were dead and unable to respond. Please note that at 2 dpa, the melanocytes lost their original shape and appeared as black smears. Scale bars are 2 mm.

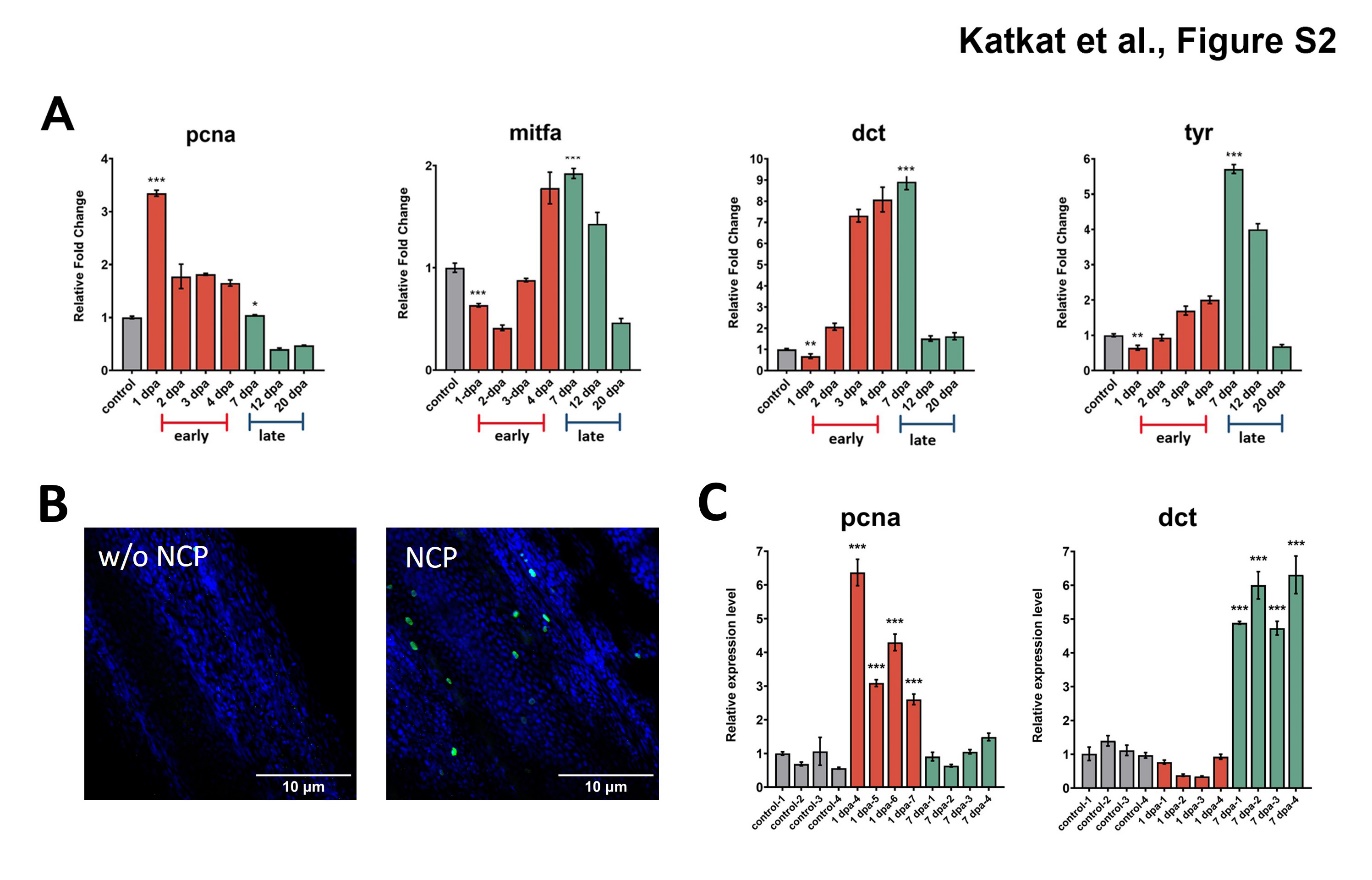

**Figure S2.** **Determination of early and late stages of melanocyte regeneration by qPCR (A)** qPCR validations of *pcna*, *mitfa*, *dct* and *tyr* in control and NCP-treated pooled fin samples at 1 dpa, 2 dpa, 3 dpa, 4 dpa, 7 dpa, 12 dpa and 20 dpa. **(B)**. Anti-phospho-Histone H3 (green) and anti-PCNA (red) staining of adult zebrafish fin that is not treated (w/o NCP) and treated with NCP for 1 day. **(C)** qPCR validations of *pcna* and *dct* in individual control and NCP-treated individual fin samples at 1 dpa and 7 dpa. *rpl13* was used as the housekeeping control gene. Error bars represent ± standard error of mean (SEM, n=3). Statistical significance was evaluated using unpaired t-test. *p<0.05, **p<0.01 and ***p<0.001.

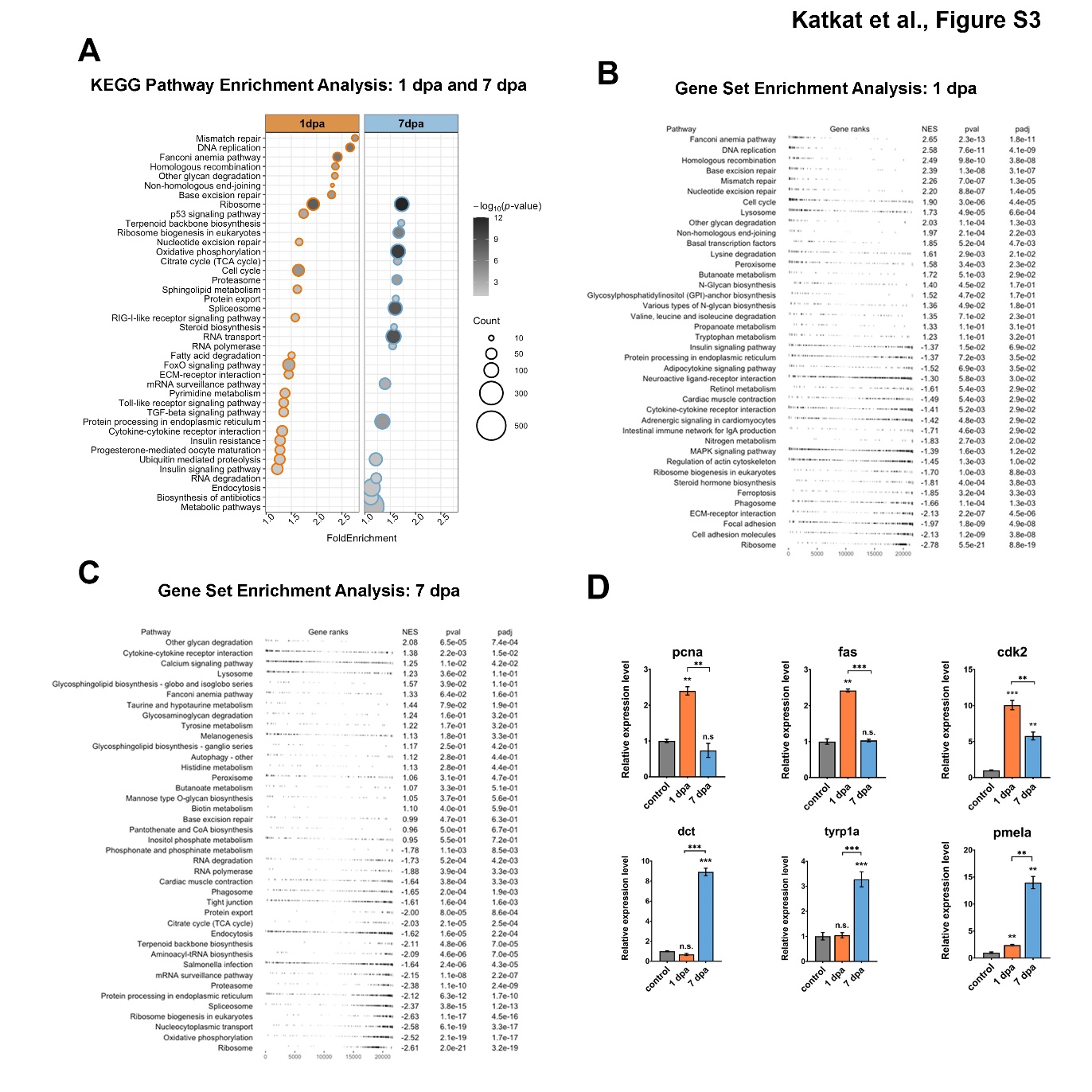

**Figure S3. KEGG pathways and GSEA enriched at two stages of the melanocyte regeneration determined by using all DEGs. (A)** DAVID was used to show the most significantly enriched KEGG pathways based on transcriptional changes at 1 dpa and 7 dpa of melanocyte regeneration. All DEGs were used for the analysis. The heatmap’s scale shows -log_10_ of EASE *p*-values for all significantly enriched GO terms. GSEA of differential expression profile of zebrafish **(B)** 1 dpa and **(C)** 7 dpa samples. Top 20 up and downregulated gene sets enriched with gene ranks, normalized enrichment score (NES), p-value, and FDR q-value. dpa: days post-ablation, DAVID: Database for Annotation, Visualization, and Integrated Discovery, KEGG: Kyoto Encyclopedia of Genes and Genomes. GSEA: Gene Set Enrichment Analysis (GSEA). **(D)** qPCR validations on 1 dpa and 7 dpa samples for *pcna*, *fas*, *cdk*, *dct*, *tyrp1a,* and *pmela*. *rpl13* was used as the housekeeping control gene. Error bars represent ± standard error of the mean (SEM, n=3). Statistical significance was evaluated using an unpaired t-test. **p<0.01, ***p<0.001 and ns: non-significant.

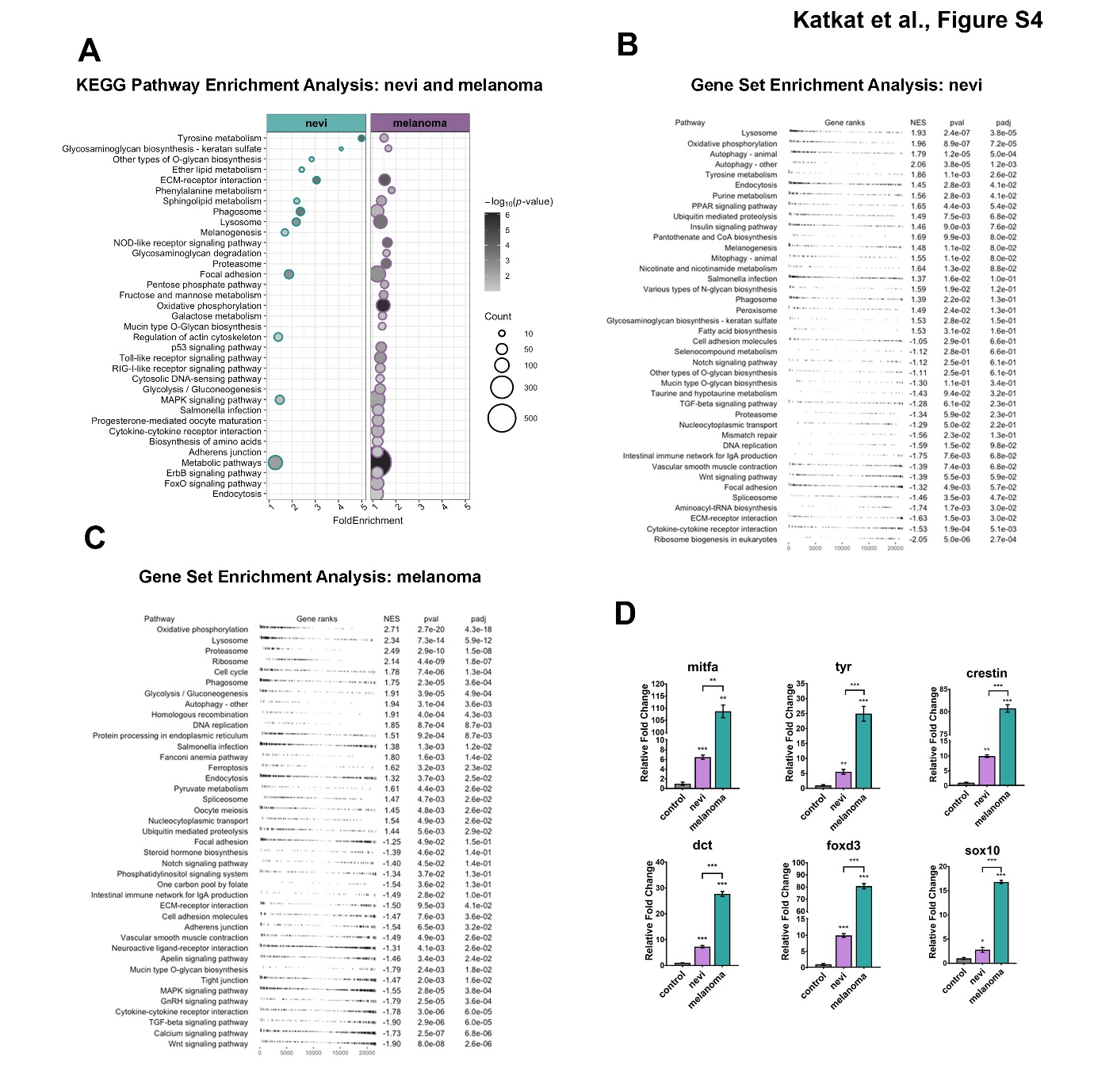

**Figure S4. KEGG pathways and GSEA enriched in nevi and melanoma determined by using all DEGs. (A)** DAVID was used to show the most significantly enriched KEGG pathways based on transcriptional changes at nevi and melanoma. All DEGs were used for the analysis. The heatmap’s scale shows -log_10_ of EASE *p*-values for all significantly enriched GO terms. GSEA of differential expression profile of zebrafish **(B)** nevi and **(C)** melanoma samples. Top 20 up and downregulated gene sets enriched with gene ranks, normalized enrichment score (NES), p-value, and FDR q-value. dpa: days post-ablation, DAVID: Database for Annotation, Visualization, and Integrated Discovery, KEGG: Kyoto Encyclopedia of Genes and Genomes. GSEA: Gene Set Enrichment Analysis (GSEA). **(D)** qPCR validations on nevi and melanoma samples for *mitfa*, *tyr*, *dct*, *foxd3*, *restin,* and *sox10*. *rpl13* was used as the housekeeping control gene. Error bars represent ± standard error of the mean (SEM, n=3). Statistical significance was evaluated using an unpaired t-test. **p<0.01 and ***p<0.001.

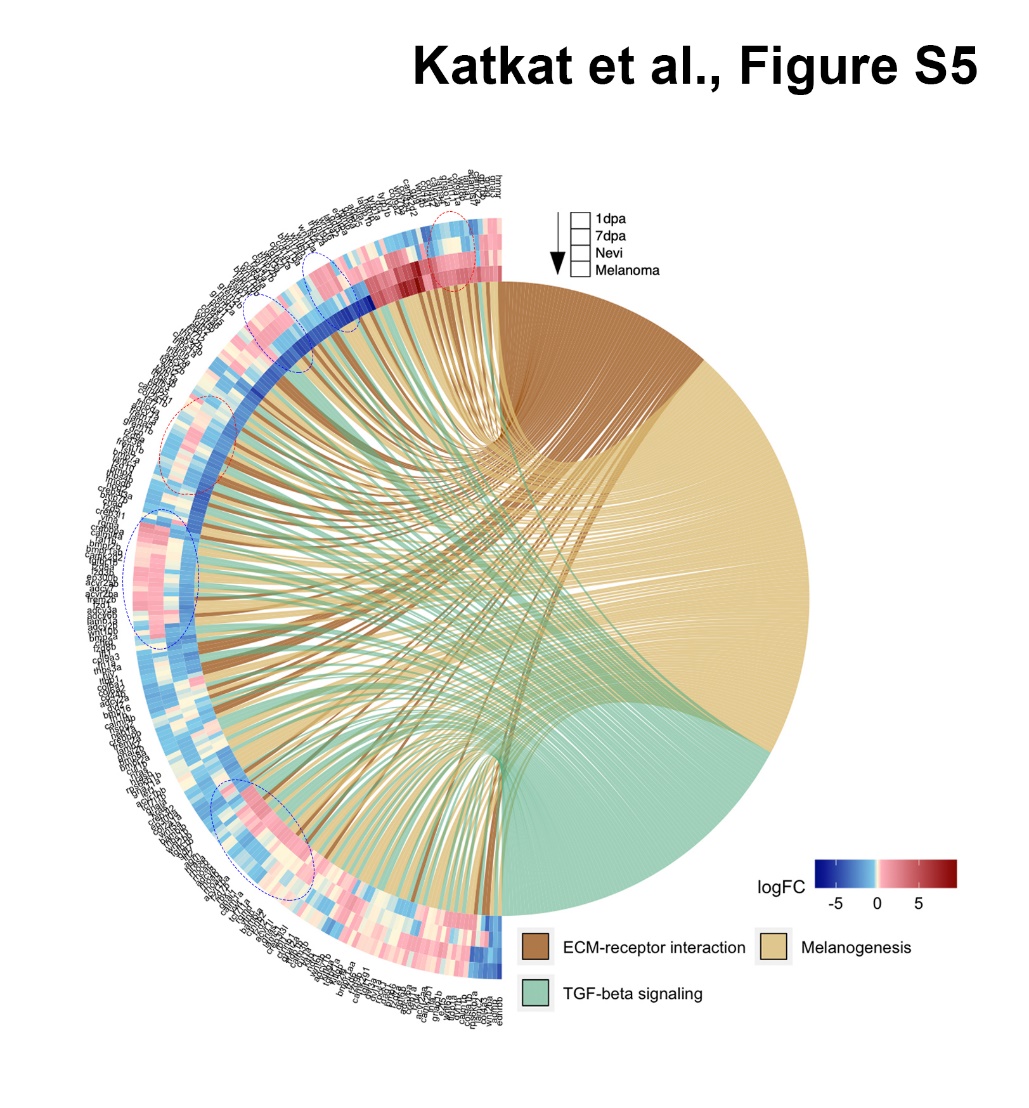

**Figure S5. Comparison of 1 dpa, 7 dpa, nevi and melanoma in terms of selected KEGG pathways.** GOChord plot shows log_2_ fold changes of the genes annotated in selected KEGG pathways for two stages of the melanocyte regeneration, nevi, and melanoma. The genes are linked to their assigned pathways by ribbons and ordered according to their log_2_ fold change values from low to high regulation, represented by a color gradient from blue to red, respectively. log_2_ fold changes are shown from the outer to the inner annulus in the following order: 1 dpa, 7 dpa, nevi and melanoma. Blue dotted circles show clusters of DEGs that are oppositely regulated between 1 dpa/7 dpa and nevi/melanoma. Red dotted circles show clusters of DEGs that are oppositely regulated between 7 dpa and nevi/melanoma. dpa: days post ablation, KEGG: Kyoto Encyclopedia of Genes and Genomes.

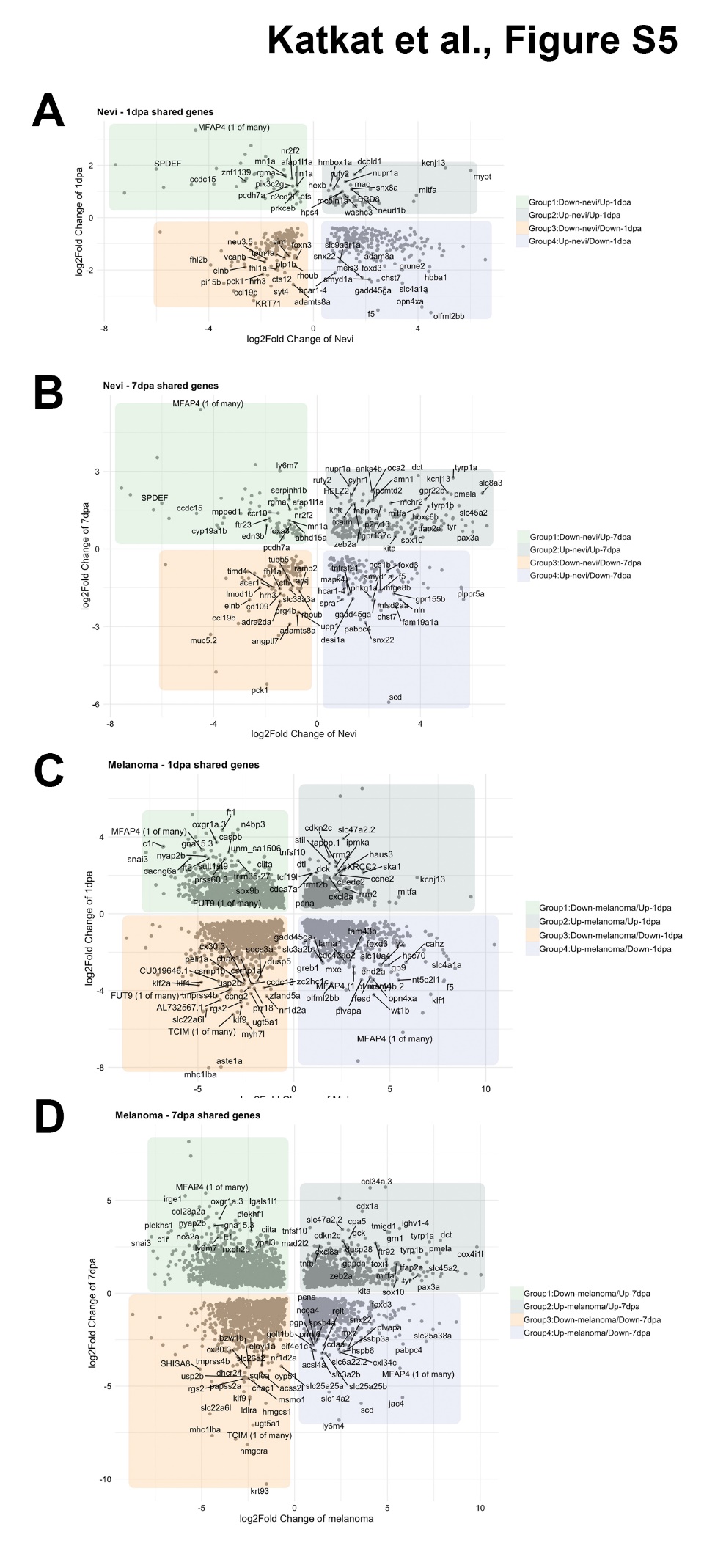

**Figure S6. Identification of shared DEGs in two stages of melanocyte regeneration, nevi and melanoma.** The 4-direction scatter plot shows differentially expressed genes (DEGs) shared between **(A)** nevi and 1 dpa, **(B)** nevi and 7 dpa, **(C)** melanoma and 1 dpa, and **(D)** melanoma and 7 dpa. Only significantly changed genes were included (Table S2). Shared DEGs were split into 4 groups based on their log fold change between conditions: Group1 genes were downregulated in nevi/melanoma but upregulated in 1 dpa/7 dpa (light green). Group2 genes were upregulated in nevi/melanoma but downregulated in 1 dpa/7 dpa (light blue). Group3 genes were downregulated in both nevi/melanoma and in 1 dpa/7 dpa (light orange). Group4 genes were upregulated in both nevi/melanoma and in 1 dpa/7 dpa (light purple).

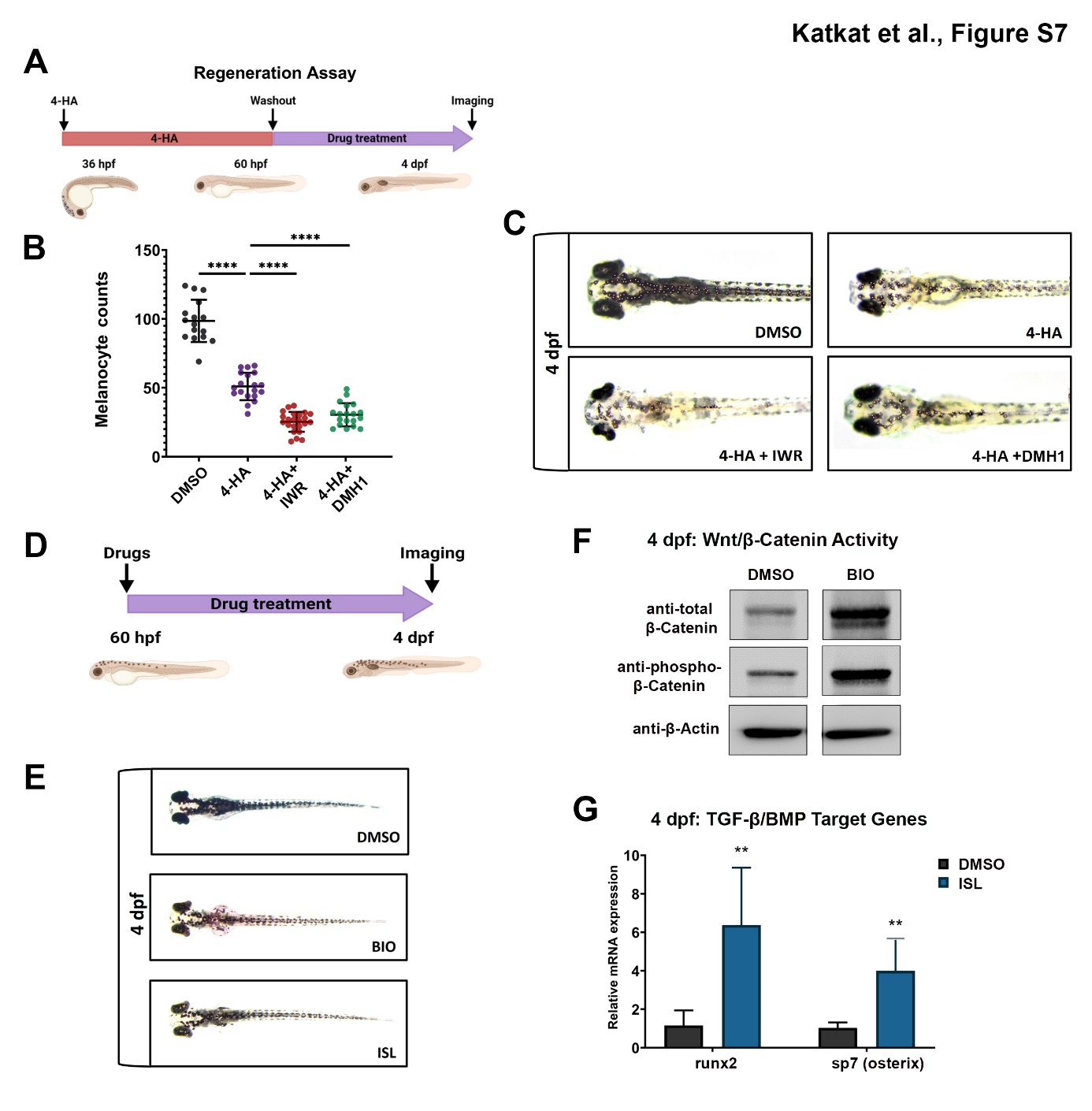
 **Figure S7. Inhibition of canonical Wnt or TGF-β/BMP signaling pathways suppresses larval melanocyte regeneration (A)** Scheme for experimental design of melanocyte regeneration. Zebrafish embryos were treated with 4-HA from 36 hpf to 60 hpf. After washout of 4-HA at 60 hpf, drugs were administered into the embryo water, and larvae were analyzed at 4 dpf. **(B)** Dot plot showing the number of melanocytes in 4 dpf larvae treated with DMSO, 4-HA, 4-HA+IWR or 4-HA+DMH1. Each dot represents one larva (DMSO n=16, 4-HA n=19, 4-HA+IWR n=23, 4-HA+DMH1 n=19). **(C)** Representative images of melanocyte regeneration groups are counted in (B). Statistical significance was evaluated using one-way ANOVA test ****p < 0.0001. **(D)** Scheme for drug treatment for zebrafish larvae. BIO or ISL was added to the embryo water at 60 hpf, and larvae were analyzed at 4 dpf. **(E)** Zebrafish larvae treated with BIO and ISL between 60 hpf-4 dpf showing no alteration in the number or pattern of melanocytes. **(F)** Western blot of zebrafish larvae treated with DMSO or ISL for total and phospho-β-catenin. **(G)** qPCR on zebrafish larvae treated with DMSO or ISL for *runx2* and *sp7 (osterix)*. *rpl13* was used as the housekeeping control gene. Error bars represent ± standard error of the mean (SEM, n=3). Statistical significance was evaluated using an unpaired t-test. **p<0.01.

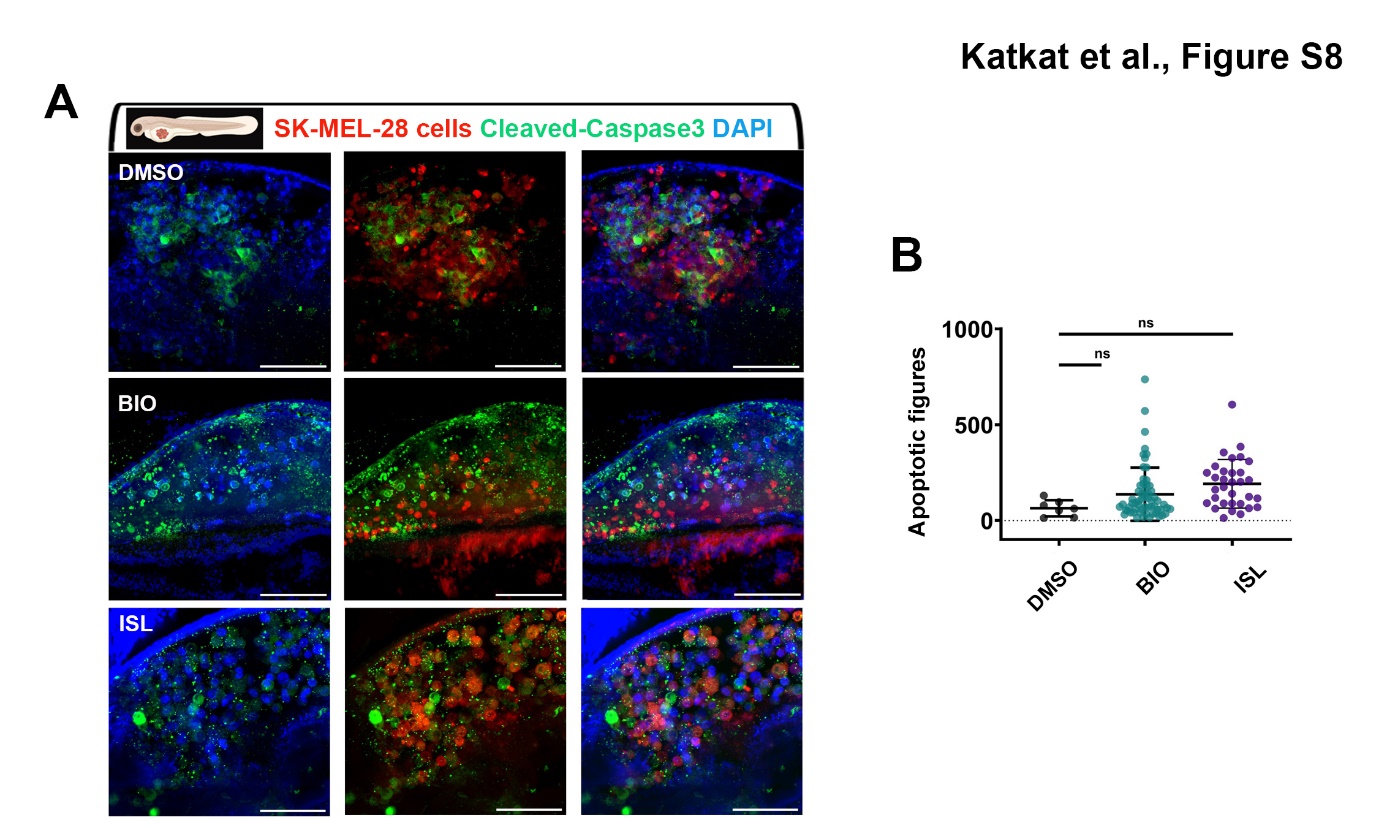
 **Figure S8. Activation of canonical Wnt or TGF-β/BMP signaling pathways does not significantly alter apoptosis of human melanoma cells *in vivo* (A)** Representative confocal microscope images of anti-cleaved-caspase-3 (green) staining of 7 dpf zebrafish larvae xenografted with SK-MEL-28 cells (red) at 2 dpf, treated with DMSO, BIO or ISL. **(B)** Dot plot showing the number of apoptotic figures of each treatment. Each dot represents apoptotic figures counted in each z-stack slice divided by the number of DiO+, DAPI+ nuclei in (A). Larvae were counterstained for DAPI. Scale bars 50μm.
